# A theory of joint attractor dynamics in the hippocampus and the entorhinal cortex accounts for artificial hippocampal remapping and individual grid cell field-to-field variability

**DOI:** 10.1101/2020.03.02.974253

**Authors:** Haggai Agmon, Yoram Burak

## Abstract

The representation of position in the brain is distributed across multiple neural populations. Grid cell modules in the medial entorhinal cortex (MEC) express activity patterns that span a low-dimensional manifold which remains stable across different environments. In contrast, the activity patterns of hippocampal place cells span distinct low-dimensional manifolds in different environments. It is unknown how these multiple representations of position are coordinated. Here we develop a theory of joint attractor dynamics in the hippocampus and the MEC. We show that the system exhibits a coordinated, joint representation of position across multiple environments, consistent with global remapping in place cells and grid cells. We then show that our model accounts for recent experimental observations that lack a mechanistic explanation: variability in the firing rate of single grid cells across firing fields, and artificial remapping of place cells under depolarization, but not under hyperpolarization, of layer II stellate cells of the MEC.

## INTRODUCTION

The discoveries of spatially selective cells in the hippocampus (O’Keefe and J., 1971) and the medial entorhinal cortex (MEC) (Hafting et al., 2005), have opened a window into the mechanisms underlying neural coding and processing of a high-level cognitive variable, which is separated from direct sensory inputs: the brain’s estimate of an animal’s position. Experimental evidence and theoretical reasoning have suggested that the neural representation of this variable is maintained by multiple attractor networks in the hippocampus (Battaglia and Treves, 1998; Quirk et al., 1990; Samsonovich and McNaughton, 1997; Schlesiger et al., 2018; Wills et al., 2005) and the MEC (Burak, 2014; Burak and Fiete, 2009; Fuhs and Touretzky, 2006; Guanella et al., 2007; McNaughton et al., 2006; Stensola et al., 2012), each expressing highly restricted patterns of neural activity at the population level that are preserved across multiple environments and conditions (Gardner et al., 2019; Trettel et al., 2019; Yoon et al., 2013). Under this interpretation of the experimental observations, patterns of population activity in the hippocampal-entorhinal system are selected from a limited repertoire, shaped by synaptic connectivity that enforces the attractor dynamics. These activity patterns are dynamically linked with the animal’s position in response to external sensory inputs and self-motion cues.

Despite the hypothesized role of recurrent connectivity in shaping spatial responses in the hippocampus and the MEC, theoretical research on the relationship between these brain areas has mostly treated them as successive stages in a processing hierarchy involving feed-forward connectivity from grid cells to place cells, or vice-versa, (Fig. 1a) [but see (Laptev and Burgess, 2019; Rennó-Costa and Tort, 2017)]. It is easy to show that summed inputs from multiple grid cells, combined with thresholding or competitive dynamics, can produce place-cell like responses (de Almeida et al., 2009; Monaco and Abbott, 2011; Neher et al., 2017; Rolls et al., 2006; Solstad et al., 2006). It is also straightforward to produce grid-cell like responses by combining synaptic inputs from multiple place cells. Moreover, theoretical studies have demonstrated that plasticity mechanisms, acting between place cells and their post-synaptic partners in the MEC, can give rise to synaptic connectivity that produces grid-cell like spatial tuning (D’Albis and Kempter, 2017; Dordek et al., 2016; Kropff and Treves, 2008; Monsalve-Mercado and Leibold, 2017; Stepanyuk, 2015; Weber and Sprekeler, 2018). Nevertheless, the functional relationship between grid cells and place cells, its relationship with recurrent connectivity within each brain region, and its role in the encoding of positions across multiple environments, remain unclear.

**Fig. 1:**
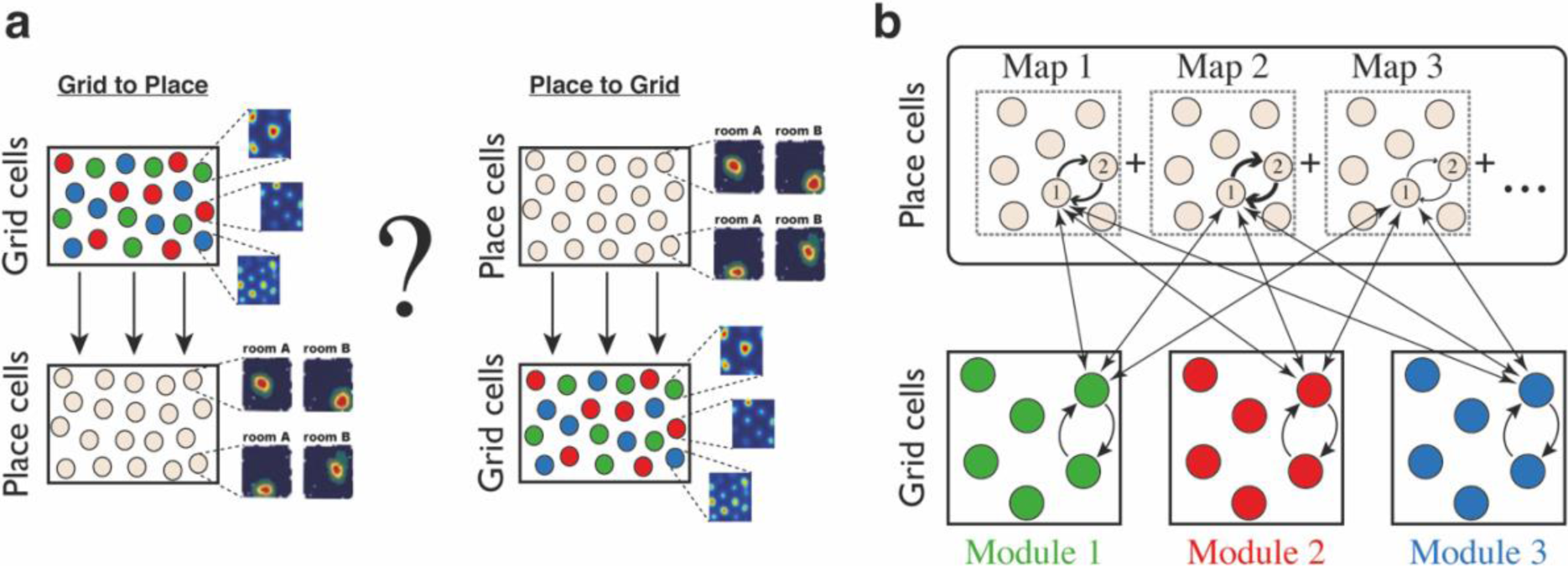
Schematic illustration: possible architectures of grid cell and place cell connectivity. **a**, Two hypotheses on connectivity, with hierarchical relationship between grid cells and place cells. Left: feed-forward connectivity from grid cells to place cells. Right: feed-forward connectivity from place cells to grid cells. Grid cells are color coded by their allocation to modules. Place cells exhibit global remapping between distinct environments (room A/room B) accompanied by grid cells realignment (not shown). **b**, Schematic illustration of the model’s architecture: place cells and grid cells are bidirectionally coupled. Top: the strength of synaptic connectivity between a pair of place cells is a sum of map-specific contributions arising from the distinct maps. In each discrete map, the connectivity depends on the environment-specific spatial separation between the place cell receptive fields (arrow thicknesses for place cells #1 and #2). Bottom: grid cell modules are modeled as independent continuous attractors. Recurrent connectivity is identical in all modules. Straight arrows: the bi-directional connection between a grid cell and a place cell is proportional to the overlap between their receptive fields across the discrete maps.

There are several difficulties with the hypothesis that spatial selectivity in the entorhinal-hippocampal system emerges from a simple feedforward architecture. First, a number of experiments indicate that grid cell inputs are not necessary for emergence of place cell spatial specificity. Stable place fields are expressed in the hippocampus of preweanling rats before the appearance of stable grid cell activity in the MEC (Langston et al., 2010; Wills et al., 2010), and place fields are maintained under MEC lesions (Hales et al., 2014). Global hippocampal remapping can occur without input from the MEC (Schlesiger et al., 2018), and place cells are expressed under the inactivation of the medial septum, which eliminates the spatial periodicity of grid cells (Koenig et al., 2011). Furthermore, new place fields can be established in novel environments under medial septum inactivation (Brandon et al., 2014).

Second, there are difficulties with the opposite hypothesis, that feedforward inputs from place cells are the sole determinants of grid cell specificity. One such difficulty arises from the observation that place cell activity patterns remap in different environments (Muller and Kubie, 1987). If grid cell responses emerge under activity dependent plasticity of feedforward synapses from place cells to grid cells (D’Albis and Kempter, 2017; Dordek et al., 2016; Kropff and Treves, 2008; Monsalve-Mercado and Leibold, 2017; Stepanyuk, 2015; Weber and Sprekeler, 2018), it is difficult to explain why they remain grid-like across familiar and novel environments, and why phase relationships between grid cells within a module are tightly preserved (Gardner et al., 2019; Trettel et al., 2019; Yoon et al., 2013). Furthermore, grid cells maintain their correlation structure under hippocampal inactivation (Almog et al., 2019), indicating that the low dimensionality of activity within each grid cell module is independent of hippocampal inputs.

Third, anatomically, the MEC and the hippocampus are reciprocally connected (Van Strien et al., 2009). A significant proportion of excitatory projections to place cells that arise in MEC originate in grid cells (Zhang et al., 2013), and there are abundant feedback projections from place cells to deep layers of MEC (Deadwyler et al., 1975).

Here, instead of assuming a feed-forward architecture of the entorhinal-hippocampal network, we explore theoretically an alternative hypothesis, that grid cells and place cells are driven by two types of synaptic inputs (Fig. 1b): inputs arising from recurrent connectivity within each brain area that restrict activity to lie within low dimensional manifolds, and inputs arising from reciprocal connections between the MEC and the hippocampus that coordinate their joint representation of position.

We first examine whether such an architecture can enforce a coordinated representation of position in both brain regions. The joint system must be able to represent all possible positions within each environment by coordinated and persistent patterns of neural activity. To achieve this goal, while accounting for the global remapping of place cells (Muller and Kubie, 1987) and the phase shift of grid cells (Fyhn et al., 2007) across distinct environments (Jeffery, 2011), we evoke theoretical ideas on the representation of multiple spatial maps by attractor dynamics in the hippocampus (Battaglia and Treves, 1998; Monasson and Rosay, 2013; Samsonovich and McNaughton, 1997), and extend them to construct synaptic connectivity that supports a representation of position jointly with the grid cell system.

We demonstrate that the coupling between hippocampal and entorhinal attractor networks enables two computational functions. First, integration of velocity inputs in the grid cell network drives similar updates of the place cell representation of position. Idiothetic path integration can thus be implemented within the grid cell system, using fixed synaptic connectivity that does not require readjustment in new environments. Second, the reciprocal connectivity between grid cells and place cells yields an error correcting mechanism that eliminates incompatible drifts in the positions represented by distinct grid cell modules. Such independent drifts would otherwise rapidly lead to catastrophic readout errors of the grid cell representation (Burak, 2014; Fiete et al., 2008; Mosheiff and Burak, 2019; Sreenivasan and Fiete, 2011; Welinder et al., 2008), and would therefore be highly detrimental for the coding of position by grid cell activity.

Next, we show that our model exhibits two emergent features that provide a compelling explanation for several recent experimental observations.

### Variability of individual grid cell firing rates across firing fields

Even though grid cell activity is spatially periodic to a good approximation, several recent works showed that firing rates of individual grid cells vary across firing fields (Dunn et al., 2017; Ismakov et al., 2017). These firing rate differences are retained across multiple exposures to the same environment. On their own, attractor models of grid cell dynamics predict that firing rates should be identical across firing fields. However, external inputs to the grid cell system could modulate the activity of grid cells in a non-periodic manner. Such inputs could have a sensory origin and project to grid cells from the LEC or the hippocampus. Here we show that even without any sensory inputs projecting into the system, the embedding of multiple spatial maps within the hippocampus, combined with the synaptic projections from place cell to grid cells, generates variability in the firing rate of grid cells across firing fields.

### Artificial hippocampal remapping upon grid cell depolarization

Place cell firing fields were recently recorded during hyperpolarization or depolarization of layer II stellate cells in the MEC (Kanter et al., 2017), giving rise to surprising observations. First, depolarization in the MEC led to an unusual form of remapping observed in hippocampal spatial responses, which differs from classical global remapping. Some place cells shifted their firing fields to new locations, while other place fields remained anchored to their original positions, and exhibited only rate remapping. This implies that in contrast to global remapping, population patterns of activity under depolarization of the MEC retain partial overlap with firing patterns that were expressed under non-perturbed conditions. We refer to this form of remapping induced by MEC depolarization as *artificial remapping*, following (Kanter et al., 2017). Second, an asymmetry was observed between consequences of depolarization and hyperpolarization in the MEC, since hyperpolarization had only mild effect on the spatial tuning curves of place cells. So far, these observations have lacked a satisfying theoretical explanation. We show below that our model exhibits the same phenomenology as observed in (Kanter et al., 2017). The theory predicts that MEC depolarization, but not hyperpolarization, alters the manifold of persistent population activity patterns, such that the hippocampus expresses mixtures of patterns that were associated with different environments before the perturbation. The emergence of such mixed states under certain perturbations is characteristic of models of associative memory. Thus, artificial remapping may offer the first experimental observation of this phenomenon in a well characterized neural network.

## RESULTS

We first ask whether bidirectional connectivity between grid cells and place cells can enforce persistent states of activity that enable continuous coding of position, and are compatible with the observed phenomenology of the place and grid cell neural codes in multiple environments: global hippocampal remapping (Muller and Kubie, 1987), and phase shifts in individual modules of the MEC (Fyhn et al., 2007). For simplicity, and to reduce the computational cost of simulations, we consider throughout this manuscript a one-dimensional (1d) analogue of grid cell and place cell dynamics.

### Model architecture and description

We model neural activity using a standard rate model. Neurons are grouped into several sub-populations: hippocampal place cells and three grid-cell modules. The architecture of synaptic connections is briefly described here, and fully specified in *Methods*. There are three types of synaptic connections in the model:

#### Recurrent connectivity between grid cells

Connectivity within each grid cell module produces a continuous attractor with a periodic manifold of steady states. We adopt the simple synaptic architecture proposed in (Guanella et al., 2007). In the 1d analogue, this form of connectivity maps into the ring attractor model (Ben-Yishai et al., 1995; Redish et al., 1996; Skaggs et al., 1995; Zhang, 1996): neurons, functionally arranged on a ring, excite nearby neighbors, while global inhibitory connections imply that distant neurons inhibit each other. This synaptic architecture leads to coactivation of a localized group of neurons, which can be positioned anywhere along the ring. The possible positions of the bump are associated with the 1d spatial coordinate by tiling them periodically along the spatial dimension, with an environment specific phase, in accordance with the experimentally observed phase-shifts of grid cell responses under global remapping (Fyhn et al., 2007).

#### Recurrent connectivity between place cells

Connectivity between place cells is based on similar principles, with two differences: first, each population activity pattern is mapped to a single spatial location. Second, the network possesses a discrete set of continuous attractors, each mapping a distinct environment to a low dimensional manifold of population activity patterns. The synaptic connectivity can be expressed as a sum over contributions from different maps, in similarity to the sum of contributions associated with discrete memories in the Hopfield model (Hopfield, 1982). To mimic the features of global remapping, spatial relationships between place cells in the different maps are related to each other by a random permutation. This is similar to previous models (Battaglia and Treves, 1998; Monasson and Rosay, 2013; Samsonovich and McNaughton, 1997) but adapted here to the formalism of a dynamical rate model.

#### Connectivity between grid cells and place cells

we first define an idealized tuning curve for each grid cell and each place cell, in each environment. These tuning curves are determined by the mapping between attractor states and positions, as described above and in *Methods*. For each pair of grid and place cells we evaluate the correlation between their spatial tuning patterns across all spatial positions and environments. The efficacy of the excitatory synapse is then linearly related to this correlation coefficient.

### Coordinated persistent representations

We demonstrate that coordinated persistent representations, and only them, simultaneously co-exist in grid cells and place cells. Our methodology is based on quantitative mapping of persistent states obtained when starting from multiple types of initial conditions. We require that in all cases the system will settle on a persistent activity bump in one of the hippocampal maps, and on persistent activity bumps in each grid cell module at matching locations. Examples of such states are shown in Figure 2. Each state exhibits unimodal and co-localized activity bumps when cells are ordered according to their firing location in a particular environment, whereas scattered place cell activity accompanied with grid cell realignment is observed when cells are ordered according to their firing location in other environments.

**Fig. 2:**
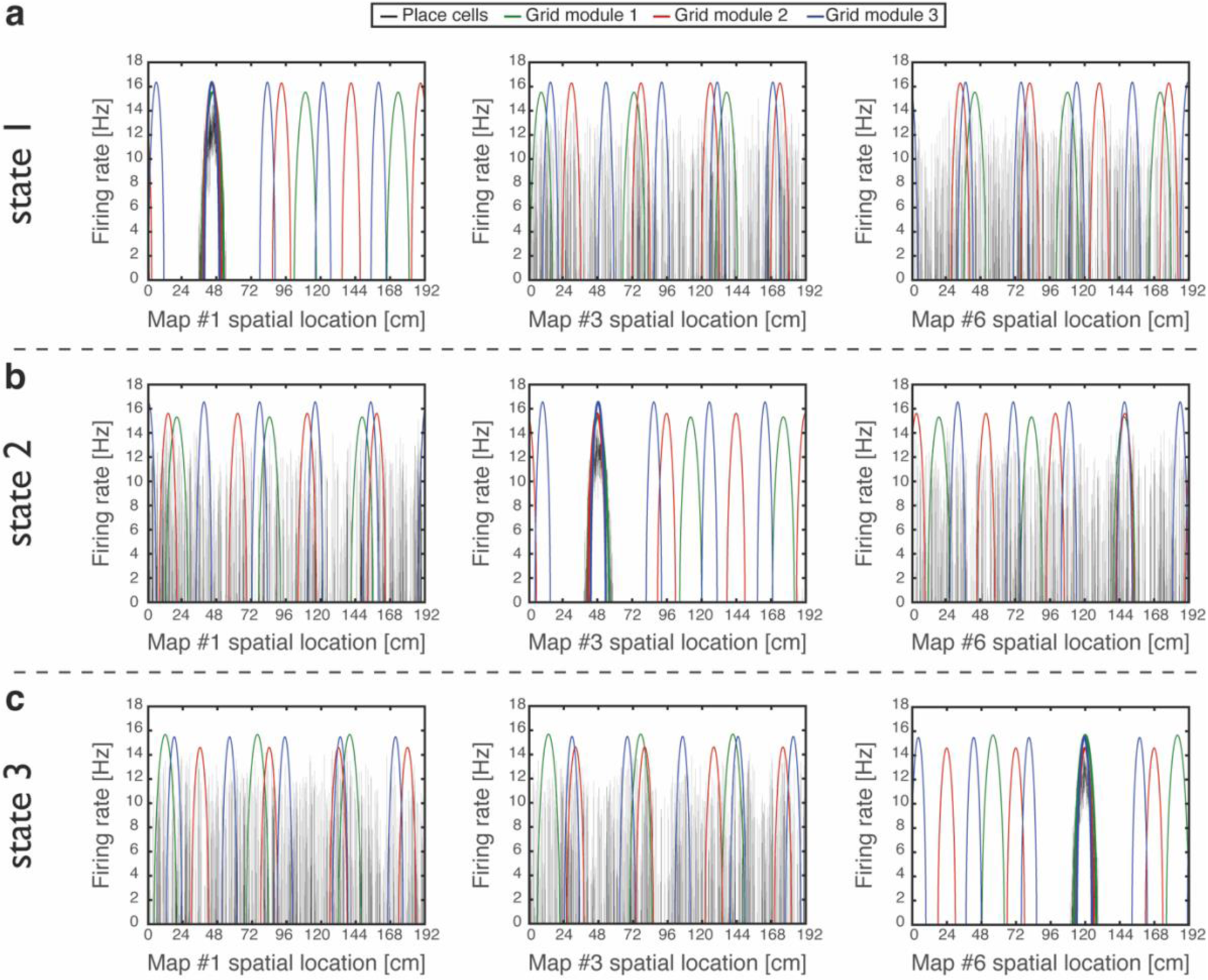
Examples of simulation results, demonstrating joint persistent states of place cells and grid cells. **a**, One persistent state of the network (state 1). In each panel, cells are ordered according to their preferred firing locations in one environment. In each grid cell module, positions displaced by any integer multiple of the grid spacing correspond to the same cell, whereas distinct place cells span the whole extent of the environment. When cells are ordered according to their preferred firing locations in map #1 (left), a unimodal place cell bump is observed, and the distinct grid cell bumps of the three modules colocalize around the same spatial location. When the cells are ordered according to their preferred firing locations in a different map (middle - map *#3* and right - map *#6)*, the same place cell activities are scattered throughout the environment and do not resemble a unimodal activity bump. Furthermore, grid cell bumps do not necessarily co-localize around the same spatial location, as they independently realign. **b-c**, Examples of persistent states (state 2 and 3), similar to (a) but representing positions in map *#3* and in map #6 respectively. Note that here the activity of place cells seems scattered and grid cells bumps independently realign when cells are ordered according to their preferred firing locations in map #1.

To test whether a persistent state represents coherently a valid position in one of the environments, we define a *bump score* that quantifies the existence of a unimodal activity bump in each sub-network. In the hippocampal sub-network, a high bump score should be seen only in a single spatial map. Within that map, we also measure the spatial location that corresponds to activity bumps in each sub-network (*Methods*). This measurement allows us to verify that all sub-networks express activity bumps in compatible positions, and to test the persistence of these bump states. Figure 3 shows the results from several types of initial conditions:

**Fig. 3:**
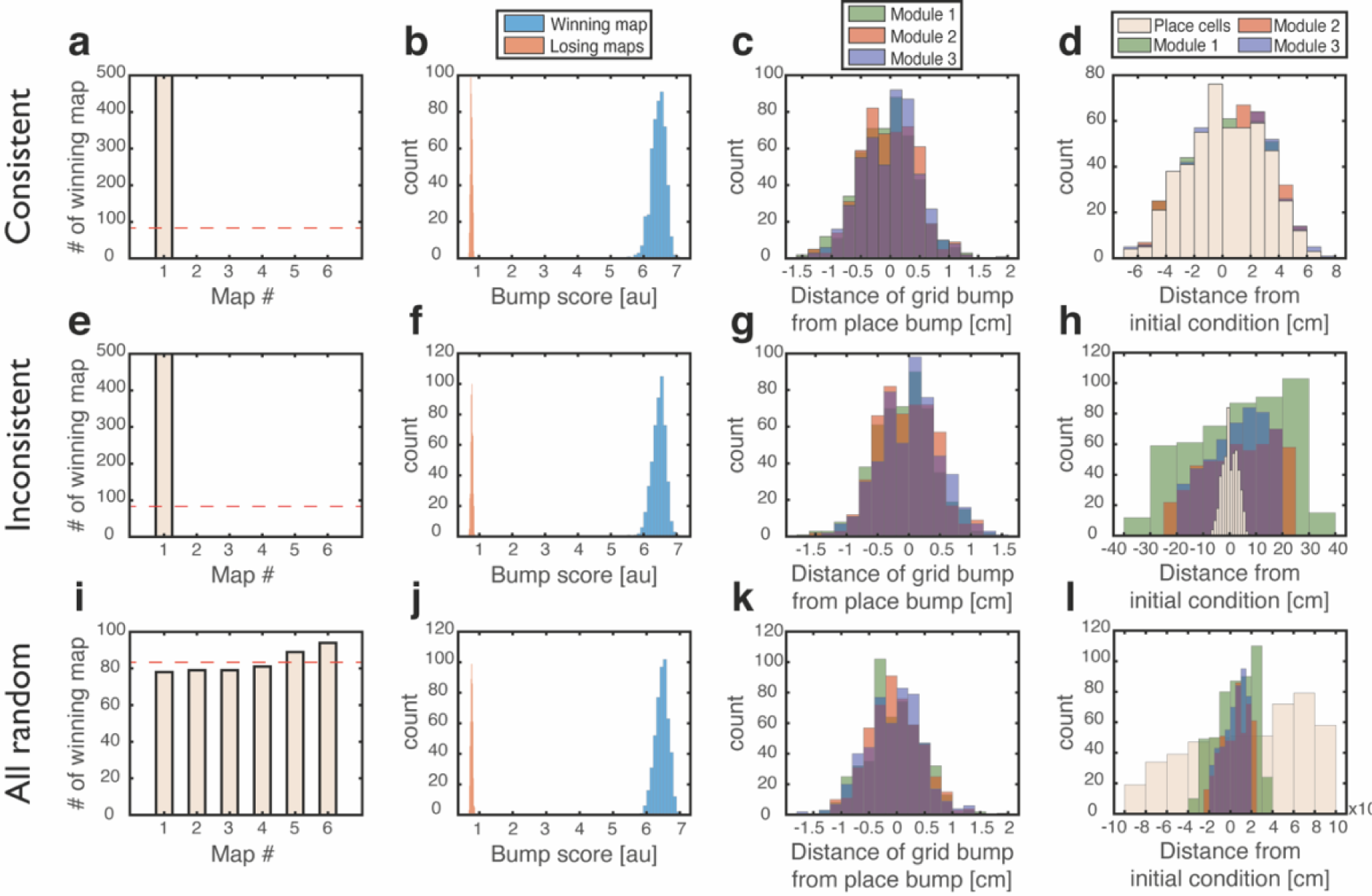
Quantitative mapping of the persistent states expressed by the joint network. In all panels, the system is placed at 500 initial conditions of three types (a-d: ‘Consistent’, e-h: ‘Inconsistent’, i-l: ‘All random’). Its state is then analyzed after a 1s delay period. **a-d**, Results from the ‘Consistent’ initial condition (the system is placed without loss of generality in map #1). **a**, Histogram of winning map, defined as the map that achieved the highest bump score. Red dashed line: uniform distribution. **b**, Histogram of place cell bump scores, obtained from the winning map (blue) and from an average on all other maps (‘Losing maps’, orange). **c**, Histogram of distances between the measured spatial location of each grid cell bump and the place cell bump, evaluated in the coordinates of the winning map. **d**, Histogram of distances traveled by the activity bumps from their initial condition, evaluated in the coordinates of the winning map. **e-h**, Similar to (a-d) but for ‘Inconsistent’ condition. **i-l**, similar to (a-d) but for ‘All random’ condition. Note that for all initial conditions the state of the system after the delay period corresponded to a single spatial map (b,f,j), and the place cell and grid cell representations are coordinated (c,g,k).

1. <u>‘Consistent condition’:</u> Grid cell and place cell activities are initially set to encode an identical location within the same map. Using these initial conditions, we verify that all locations, in all environments, can be represented by persistent states (Fig. 3a-d). The bump score remains high after 1s when initializing the network in co-localized bump states, and the drift of the represented position is small compared to the width of activity bumps, indicating that these states are persistent.
2. <u>‘Inconsistent condition’:</u> Grid cell and place cell activities are initially set to encode different locations. Using this test, we verify that such incompatible states are unstable and give rise (after a short transient period) to a consistent state (Fig. 3e-h).
3. <u>‘All random’:</u> Both grid cell and place cell activities are set to random initial rates. The goal of these tests is to verify that even if bump states do not exist initially, the system evolves into unimodal activity bumps in compatible positions, and that no other spurious states emerge. As expected, co-localized place cell and grid cell activity bumps appear with equal probabilities across the different maps (Fig. 3i).
4. Finally, we test how the system evolves from two additional types of initial states: in one condition grid cells are initially set to have random rates, while place cells are set to encode a specific location. In the second condition, grid cells are set to encode a specific location, and place cells are assigned with random rates. The results (Supplementary Fig. 1) indicate that both place cells and grid cells are capable of attracting activity in the other network to organize in a compatible manner with their initial state.

### Functional consequences: path integration and dynamic coupling

Two functional consequences arise from the coupling between the place and grid cell networks:

#### Path integration

The MEC has been hypothesized to be the brain area responsible for idiothetic path integration for several reasons (Burak, 2014; Hafting et al., 2005; McNaughton et al., 2006) among them the highly geometric nature of grid cell spatial selectivity, the existence of inputs to the MEC that convey information about the animal’s self-motion and heading (Kropff et al., 2015; Sargolini et al., 2006), and the impairment of path integration following disruption of grid cell activity (Gil et al., 2018). The similarity in response patterns of grid cells across environments implies that implementation of path integration in the MEC could be accomplished in different environments by the same synaptic connectivity, whereas implementation in the hippocampus (McNaughton et al., 1996) would require establishment of dedicated synaptic connectivity in each environment. Figure 4a demonstrates that in the coupled system, integration of velocity inputs in the MEC, using mechanisms described in (Burak and Fiete, 2009) (*Methods*) induces coordinated updates in the position represented in the place cell network. Figure 4b shows that errors accrued during path integration remain small, when using realistic velocities that were measured experimentally during foraging in an open field environment (Fig. 4c). Interestingly, the representation in the hippocampus lags behind the entorhinal representation, due to synaptic transmission delays (Fig. 4d).

**Fig. 4:**
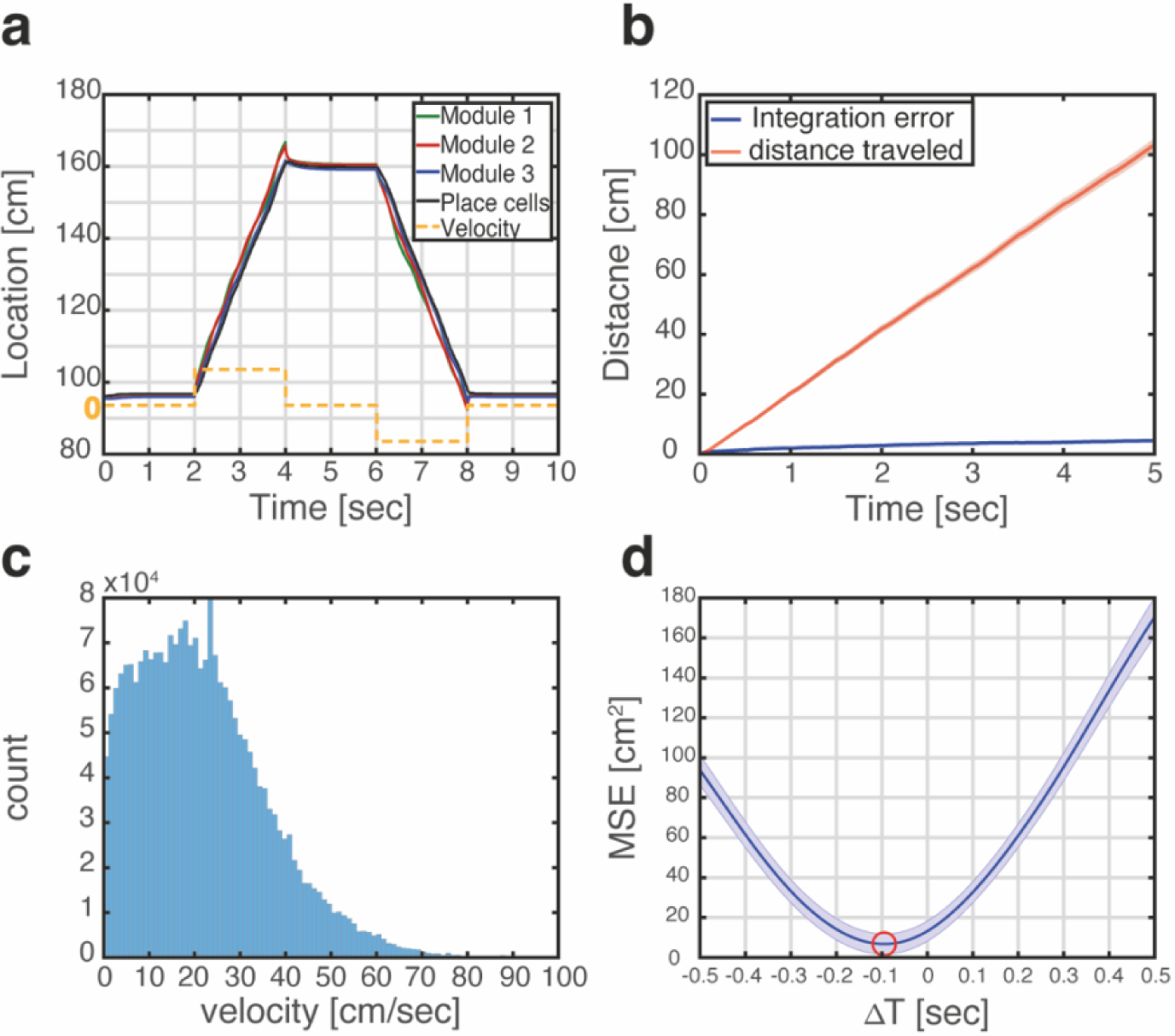
Velocity integration in grid cells can update the place cell representation. **a**, Readout of place cell bump location (black) while grid cell modules (green, red, and blue) integrate a velocity profile (dashed orange line). **b**, Mean absolute distance between measured location of the place cell activity bump, and integrated velocity (blue), as a function of time. This quantity is compared with the total distance traveled by the place cell bump (red), which is much larger. Averaging was performed over 500 trajectories. In each trajectory velocity was randomly sampled every 0.5sec from an experimentally measured velocity distribution, which was obtained during foraging of a rat in an open field environment (see panel c). Shaded error bars are 1.96 times the standard deviations obtained from each simulation, divided by the square root of the number of realizations (corresponding to a confidence interval of 95%). **c**, Histogram of velocities measured experimentally during foraging of a rat in an open field environment (Hafting et al., 2005). **d**, Mean squared displacement between positions represented by place cells and grid cells, when evaluated at varying time lags between the two measurements. Red circle marks the minimal MSE. The minimum MSE occurs at a negative time lag (∼-100 ms), which indicates that the place cell representation lags behind the grid cell representation. Shaded error bars are 1.96 times the standard deviations obtained from each simulation, divided by the square root of the number of realizations (corresponding to a confidence interval of 95%).

#### Coordination of the grid cell representation: suppression of incompatible drifts

The modular structure of the grid cell code for position confers it with large representational capacity (Fiete et al., 2008). Yet, the modularity poses a significant challenge for the neural circuitry that maintains the representation and updates it based on self-motion. Under conditions in which sensory inputs are absent or poor, incompatible drifts can accrue in the phases of different modules. Such incompatible drifts rapidly lead to combinations of spatial phases that do not represent any position in the nearby vicinity of the animal, resulting in abrupt shifts in the represented position (Fig. 5a) [see also (Burak, 2014)]. The difficulty arising from occurrence of such catastrophic readout errors has been identified in early works on grid cell coding (Fiete et al., 2008). Since then, two possible solutions to this problem were proposed. In one solution (Sreenivasan and Fiete, 2011; Welinder et al., 2008) the hippocampal network reads out the position jointly represented by grid cell modules, and projections from hippocampus to the MEC correct small incompatible drifts accrued in the different modules. A second solution, proposed more recently (Mosheiff and Burak, 2019) involves synaptic connectivity between grid cells belonging to different modules. The first solution has been explored previously (Sreenivasan and Fiete, 2011) without explicit modelling of attractor dynamics in the hippocampus, and without considering the embedding of multiple spatial maps in the entorhinal-hippocampal network. Therefore, we tested whether coordination of modules can be implemented in the coupled system, across multiple environments.

**Fig. 5:**
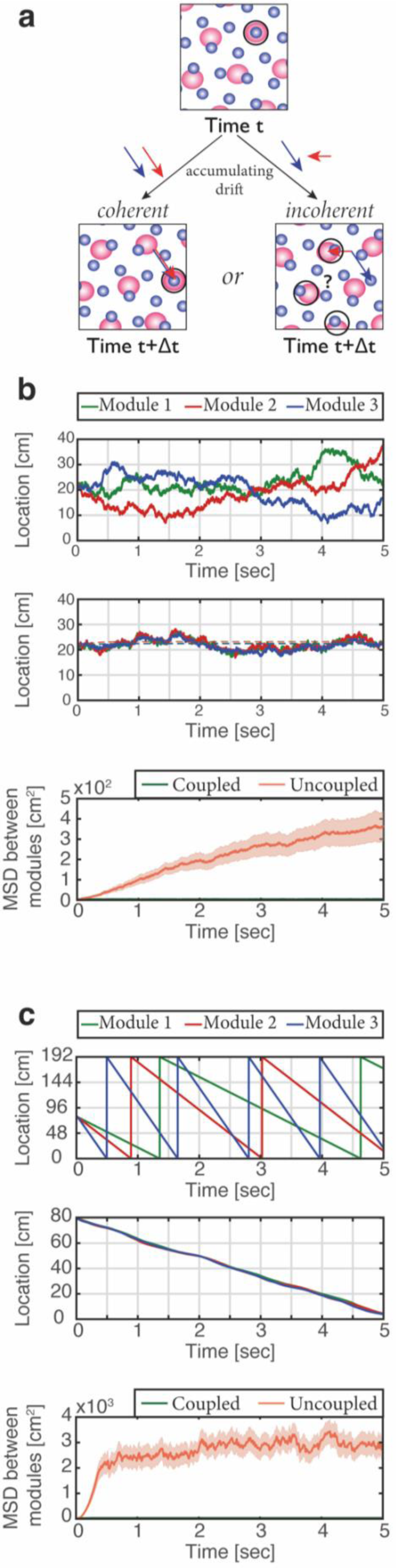
Place cells coordinate the representations of position in distinct grid cell modules. **a**, The need for coupling. Top panel: blue and red shaded areas represent schematically the posterior likelihood for the animal’s position in the environment, obtained from activity of grid cells in two different modules (each color corresponds to one module). The position which is most likely given the joint activity in both modules is designated by a black circle. Bottom left: when noise-driven drifts in the two modules are identical (coherent drifts, blue and red arrows), the outcome is a small shift in the represented position. Bottom right: incoherent drifts in the two modules may result in abrupt jumps in the maximum-likelihood location(Burak, 2014; Mosheiff and Burak, 2019). **b**, Examples of simulation results starting from a consistent initial condition showing grid cell module bump locations (green, red and blue) vs. time without (up) and with (middle) coupling between place cells and grid cells. Random accumulated drifts are driven by intrinsic neural noise, modeled as arising from Poisson spiking. Dashed lines in middle panel are generated from a non-noisy simulation for reference. Bottom: mean square displacement (MSD) between bump locations of distinct grid cell modules (averaged across all pairs and realizations of the stochastic dynamics) vs. time with (green) and without (red) coupling between place cells and grid cells, starting from a consistent initial condition. In the coupled case the MSD saturates and doesn’t exceed a value of 2cm^2^. Shaded error bars are 1.96 times the standard deviations obtained from each simulation, divided by the square root of the number of realizations (corresponding to a confidence interval of 95%). **c**, Similar analysis as (b) obtained from simulations in which drifts arise from inconsistent velocity inputs provided to the distinct modules. The drifts remain coordinated in the coupled case, and the MSD doesn’t exceed a value of 9cm^2^.

We first considered the consequences of stochastic neural firing, by modelling grid cells as noisy (Poisson) units. In the absence of coupling between the MEC and hippocampus the positions represented in different modules drifted randomly in an uncoordinated manner (Fig. 5b, top). With bidirectional coupling (Fig. 5b, middle) the representation of position accrued errors relative to the true position of the animal, but remained coordinated in all modules and in the hippocampus. Figure 5b (bottom) quantifies the relative drift between modules in the different conditions over multiple realizations of the dynamics, showing that drifts remain coordinated under coupling with the hippocampus. Drifts that are coordinated across modules are much less detrimental for decoding of position than coordinated drifts [see (Burak, 2014; Mosheiff and Burak, 2019) for a thorough discussion]. Next, we demonstrated that the system is resilient to noise in the velocity inputs to the different modules (Fig. 5c). Even in an extreme situation, in which modules received highly incompatible velocity inputs, their motion remained coordinated in the coupled network but not in the uncoupled network.

### Emergent properties in the hippocampus and MEC

The embedding of multiple spatial maps in the coupled hippocampal-entorhinal network gives rise to two non-trivial emergent properties.

#### Variability of individual grid cell firing rates across firing fields

As the animal moves in a given environment, synaptic inputs from place cells include a contribution associated with that environment, which are spatially periodic. However, grid cells also receive contributions associated with other spatial maps that are embedded in the synaptic connectivity. The latter contributions are spatially aperiodic within the present environment, leading to variability across fields. As an example, Figure 6a shows the population activity of place cells and grid cells from module #3. The system is placed at three spatial locations that are displaced from each other by the grid spacing of module #3. Thus, the same grid cells are active in these three locations. However, the amplitude of grid cell activity bumps differ at the three locations. Figure 6b quantifies the variability of peak firing rates generated by individual grid cells across firing fields within a single environment, using the coefficient of variation (CV): a histogram of the CV, collected from all cells, is shown for networks with varying number of embedded maps. CVs vanish when a single spatial map is embedded in the synaptic connectivity, and increase with the addition of spatial maps.

**Fig. 6:**
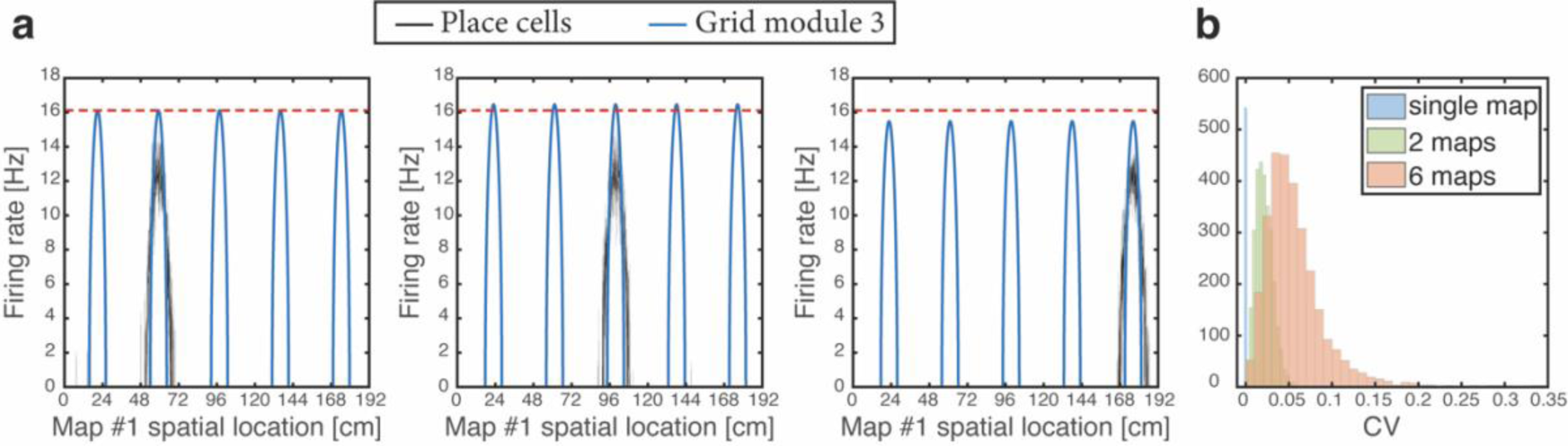
Spontaneously emerging variability of individual grid cell firing rates across firing fields. **a**, Simultaneous firing rates of place cells (black) and grid cells from one module (module #3, blue), shown at three different persistent states that represent periodic locations of module #3. Even though the same grid cells are active in all three locations, the amplitude of the grid cell activity bumps differ (compared with the red dashed line which shows the maximal grid cell firing rate achieved at the left example, for reference). Different place cells are active in the three examples, thus providing different quenched noise to the same grid cells in each case. **b**, Histogram showing CV of maximal firing rate obtained from all individual grid cells across firing fields within a single environment, shown for networks with a single map (blue), two maps (green), and six maps (red).

#### Artificial hippocampal remapping upon grid cell depolarization

In a recent experiment (Kanter et al., 2017) MEC layer 2 stellate cells were reversibly hyperpolarized or depolarized using chemogenetic receptors, while animals were foraging in a familiar environment. Chronic tetrode arrays implanted in the hippocampus were used to monitor CA1 firing fields during these manipulations. Hyperpolarization of the MEC had very little effect on place cell firing fields, whereas depolarization of the MEC elicited dramatic effects: a significant portion of place cells exhibited changes in the locations or rates of their firing fields, resembling those seen during global and rate remapping experiments. This effect included place cells that had been turned off or on as seen in natural remapping experiments. Firing fields of all other cells remained in their baseline positions, without significant changes. As for the grid cells, both manipulations led to changes in the firing rates but did not elicit a change in the firing locations, unlike the coherent shifts and rotations seen in distinct familiar environments (Fyhn et al., 2007).

The asymmetric effect of entorhinal depolarization and hyperpolarization was highlighted (Kanter et al., 2017) as the most puzzling experimental outcome. Indeed, under simple feedforward models of place cell emergence from grid cell inputs (de Almeida et al., 2009; Monaco and Abbott, 2011; Neher et al., 2017; Rolls et al., 2006; Solstad et al., 2006), it is difficult to envision why such an asymmetry would be observed. Overall, a theoretical framework that can coherently explain the consequences of depolarization and hyperpolarization of the MEC hasn’t been available so far.

To test the effect of grid cell depolarization or hyperpolarization in our model, we adjusted the excitability of grid cells by adding a fixed contribution to their synaptic drive, which was either excitatory or inhibitory. We first placed the system in persistent states in one map (map 1). We then hyperpolarized the grid cell network. Place cell bumps remained around their baseline spatial location, and activity in the place-cell network exhibited significant overlap only with map 1 (Fig. 7a-b). Similar results from control trials, in which no manipulation was performed, are shown in Figure 3a-b. The minor effect of grid cell hyperpolarization is not surprising, since the hippocampal network is structured such that it can sustain population activity patterns associated with each environment even without inputs from grid cells. Therefore, suppression of activity in the MEC network did not significantly alter the population activity patterns expressed by the place cell network.

**Fig. 7:**
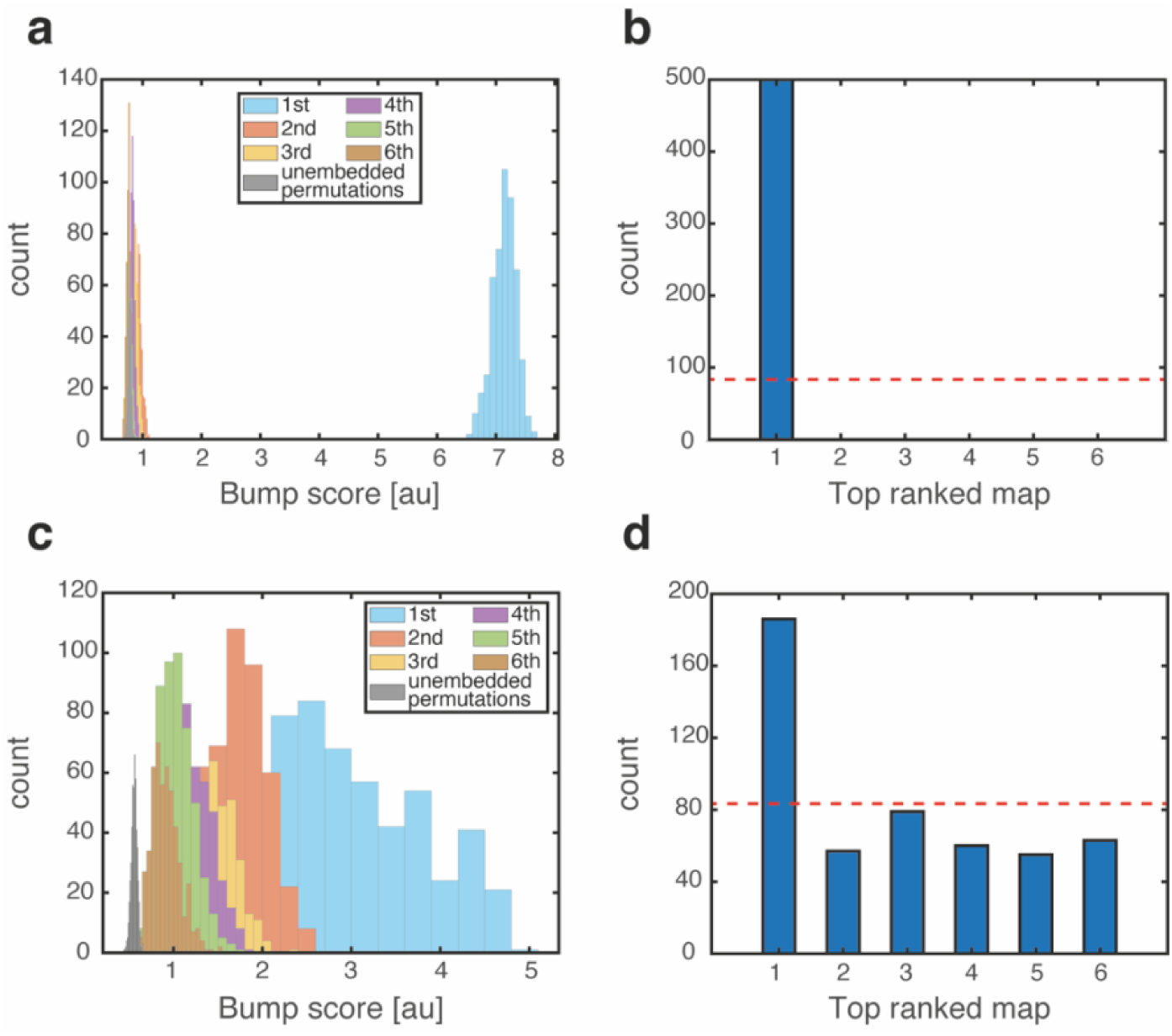
Emergence of persistent mixed states under depolarization but not under hyperpolarization of grid cells. **a**, Histogram showing bump score distributions for all embedded maps under grid cell hyperpolarization. In each simulation, the system is placed in a ‘Consistent’ initial condition, and hyperpolarization is applied. Spatial maps are then ranked according to their bump score. Each color corresponds to a histogram over all bump scores from maps with specific rank, regardless of the map’s identity (this is identical to the analysis shown in Figure 3b,g,f except that here losing maps are separated into distinct histograms according to rank). Bump scores for random unembedded maps are shown in gray. **b**, Distribution of identities of the top scored maps from (a). Dashed red line shows the uniform distribution. In all realizations, top ranked map is map #1. **c**, Same as (a) under grid cell depolarization. Grid scores are typically lower than the winning scores in Figure 3b. Bump score distributions from differently ranked maps overlap, and all histograms exhibit significantly higher bump scores than those of unembedded maps (gray), indicating simultaneous expression of activity patterns from multiple embedded maps. **d**, Same as (b) under grid cell depolarization.

In contrast with this result, depolarization of the MEC elicited significant changes in the structure of population activity patterns expressed by the hippocampal network. Following a short transient, population activity in CA1 stabilized on patterns that exhibited overlap with patterns that were previously associated with multiple spatial maps (Fig. 7c). The overlap was highly significant compared to overlaps obtained with random maps that were not embedded in the connectivity (gray bars in Fig. 7c). Thus, the place cell network expressed mixtures of population activity patterns from different maps stored in the hippocampal connectivity. Typically, the highest overlap remained with map 1 (Fig. 7d). An example of population activity patterns before and after depolarization in one location is shown in Supplementary Figure 2a-d. In similarity to experimental results (Kanter et al., 2017), both depolarization and hyperpolarization changed the firing rate, but not firing location, of MEC neurons. On average place cells increased their firing rates under grid cell depolarization and decreased their firing rates under grid cell hyperpolarization (not shown), in agreement with experiments. Supplementary Figure 2e demonstrates that the mixed activity patterns are persistent.

Our interpretation of artificial remapping is thus that excessive inputs from grid cells enhance the quenched noise arising from multiple embedded maps, and that this enhancement puts the system in a regime in which it expresses mixed states. To demonstrate the existence of such a regime, we show in Supplementary Figure 3a-b that similar expression of mixed states emerges when synapses between grid and place cell are strengthened, even without grid cell depolarization. In contrast, selective strengthening only of the component associated with one of the maps, does not generate mixed states (Supplementary Fig. 3c-d). On the contrary: this selective enhancement suppresses the emergence of mixed states under grid cell depolarization (Supplementary Fig. 3e-f). These results demonstrate that the emergence of mixed states is not simply due to enhancement of inputs from the MEC, but is due to the amplification of quenched noise.

To make direct contact with the experimental observations (Kanter et al., 2017), we examined the firing properties of individual place cells as a function of the animal’s position during locomotion under MEC depolarization. The observation that population activity patterns under depolarization are mixtures of activity patterns from different maps led us to expect that some cells would maintain place fields in their baseline locations, while others would remap. To test this hypothesis, we first placed the system in a state corresponding to a particular position in map 1. Then, while applying the perturbation, we induced path integration in the MEC network and monitored population activity patterns while continuously scanning positions along the environment.

In accordance with the hypothesis, we found that cells exhibited the same phenomenology observed experimentally (Kanter et al., 2017) (Fig. 8a-g). Many cells shifted their firing location, whereas others maintained their place fields in their baseline location, while possibly undergoing rate remapping. Some place cells expressed one or more firing fields in addition to their original field location, some turned off, and some place cells remapped into several fields. All active cells exhibited a continuous activity pattern with one, or occasionally few, unimodal firing fields. Supplementary Figure 4 shows all the population activity patterns observed while traversing the environment in a single path integration cycle. In agreement with the experimental observations, firing patterns returned to baseline configurations following reversal of MEC depolarization (not shown). As expected (Fig. 7a-b), hyperpolarization did not elicit significant changes in the firing locations of place cells (Fig. 8h).

**Fig. 8:**
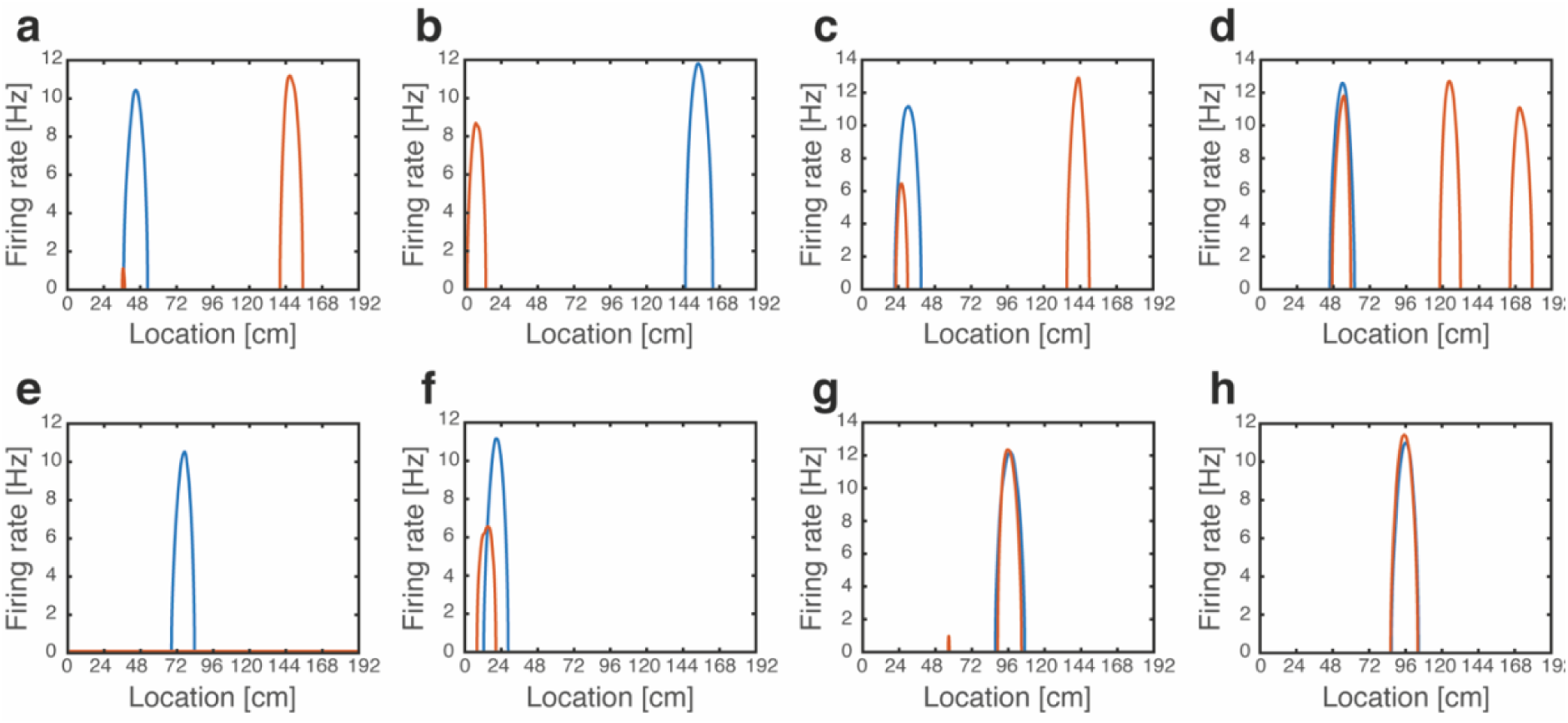
Artificial remapping of place cells following depolarization, but not hyperpolarization of grid cells. **a-g**, Firing rate maps of seven representative place cells during locomotion with (blue) and without (orange) grid cell depolarization. Rate maps show that under grid cell depolarization place cells changed locations (a-b), acquired additional fields (c-d), turned off (e), rate remapped (f) or were unaffected (g). **h,** Rate maps of a representative place cell with (blue) and without (orange) grid cell hyperpolarization.

Finally, we examined how grid cell activity is influenced by the altered firing patterns of place cells during grid cell depolarization. Our previous observation that place cell inputs induce variability in firing rates of single grid cells across different firing fields, led us to examine the rank order of grid fields according to peak firing rate before and after depolarization. Supplementary Figure 5 demonstrates that when a single spatial map is embedded in the synaptic connectivity the rank correlation is equal to unity, as expected. When multiple spatial maps are embedded in the connectivity, the rank order correlation becomes weak, in accordance with experimental observations (Kanter et al., 2017).

## DISCUSSION

We demonstrated here how multiple attractor networks in the MEC and hippocampus can be coupled to each other, such that they jointly and coherently represent positions across multiple spatial maps, while exhibiting the features of global remapping that are experimentally observed in both brain areas.

Due to the embedding of multiple spatial maps in the hippocampal network, the input to a place cell in a given environment can be separated into two components. The first component arises from the synaptic connectivity associated with the presently expressed spatial map. The other component is associated with other spatial maps, and contributes quenched noise to the synaptic inputs (Battaglia and Treves, 1998; Monasson and Rosay, 2013; Samsonovich and McNaughton, 1997) (supplementary information). Due to this quenched noise, the system loses its ability to represent all spatial positions in true steady states of the network dynamics. However, if the quenched noise is sufficiently weak, bump states representing all possible positions remain highly persistent. This feature of continuous attractor systems with quenched noise, and specifically with multiple maps, has been extensively discussed in previous works (Itskov et al., 2011; Monasson and Rosay, 2014). For this reason, we tested for the existence of persistent rather than steady states in our analysis (Fig. 3). In particular, we verified that drifts of the represented location are small, when starting from the consistent condition (Fig. 3d). The drift rate is expected to be even smaller with a realistic number of place cells, since it scales inversely with the network size (Monasson and Rosay, 2014). Furthermore, it has been demonstrated that sensory inputs, projecting to the hippocampus can pin the representation and prevent drift (Monasson and Rosay, 2014). Systematic drifts (Itskov et al., 2011; Monasson and Rosay, 2014) and pinning of the attractor state when velocity inputs are small (Burak and Fiete, 2009), can reduce the precision of path integration. Yet, Figure 4b demonstrates that with a realistic distribution of velocities, the system is still highly effective in integrating the velocity inputs, over multiple spatial maps, even when allocentric sensory inputs are completely absent.

Previous theoretical works showed that the grid cell representation has a high dynamic range, which enables coding and storage in working memory of the animal’s location with high spatial resolution, over a large range of positions (Burak, 2014; Fiete et al., 2008; Mathis et al., 2012; Sreenivasan and Fiete, 2011; Wei et al., 2015). However, due to the distributed nature of the grid cell code, it was argued that it is crucial to maintain precise coordination between the positions represented in different grid cell modules (Burak, 2014; Mosheiff and Burak, 2019; Sreenivasan and Fiete, 2011; Welinder et al., 2008). Here we showed that the required coordination can be achieved through bidirectional connectivity between grid cells and place cells, over multiple spatial maps, using an explicit model of attractor dynamics in the MEC and the hippocampus.

We showed that the embedding of multiple maps in the synaptic connectivity naturally produces variability in the firing rates of individual grid cells across different firing fields. While the experimentally observed variability may arise from place cell inputs, as predicted by our model and proposed in (Ismakov et al., 2017), it could also arise from inputs from the LEC or other grid cell modules. In either case, the variability could be driven by sensory inputs, whereas the mechanism identified in this work is independent of such inputs. The fact that the rank order of grid fields was modified under grid cell depolarization when the hippocampal network exhibited artificial remapping (Kanter et al., 2017), as observed in our model, provides further support to the hypothesis that inputs from place cells significantly contribute to the variability, independently of sensory inputs.

Our model reproduces the recently observed phenomenology of artificial remapping in the hippocampus under MEC depolarization but not under hyperpolarization. According to our model, artificial remapping is a consequence of a perturbation to the network that places it in a regime in which it expresses mixtures of population activity patterns. This transition does not require any modification of sensory inputs, in accordance with the experimental protocol (Kanter et al., 2017), in which the environment remains fixed. All types of remapping observed experimentally (Kanter et al., 2017), emerged in our model except for cells that lacked a firing field before depolarization and acquired one afterwards. This is due to the fact that, for simplicity, we assigned a spatial receptive field to all cells in each environment. It would be straightforward to include in the model silent cells in each spatial map (Monasson and Rosay, 2013), in order to obtain cells in the hippocampal network that acquire a firing field only after grid cell depolarization.

Our results also suggest that place cell artificial remapping is likely to occur at specific spatial locations where most grid cells are active across all maps, and unlikely to occur in positions in which grid cells are inactive. One manifestation of this prediction can be easily seen in Supplementary Figure 4a: a margin of positions near the diagonal into which place cells are highly unlikely to remap.

The model explains why artificial remapping occurred under depolarization, but not under hyperpolarization of MEC cells (Kanter et al., 2017). In another experiment(Miao et al., 2015) remapping in the hippocampus was observed under suppression of activity in the MEC. The reason for the different outcomes (Kanter et al., 2017; Miao et al., 2015) is unclear. The different outcomes may have arisen from specificity differences in the targeted populations: while in Ref (Kanter et al., 2017) chemogenetic receptors were targeted almost exclusively to layer II stellate cells of the MEC in transgenic animals, a broader population has been likely affected using viral injection in Ref (Miao et al., 2015).

### Experimental predictions

We next discuss some of the predictions arising from our model. To test some of these predictions experimentally, it will be necessary to simultaneously record the activity of multiple cells in the MEC and the hippocampus in freely behaving animals, in sufficient numbers that will enable dynamic decoding of the population activity. Thus, our results offer a rich set of predictions that could be tested using high-throughput recording techniques (Jun et al., 2017; Pfeiffer and Foster, 2015; Ziv et al., 2013; Zong et al., 2017).

#### Idiothetic path integration

We predict that when updates of the brain’s self-estimate of position rely solely on idiothetic path integration, the hippocampal representation of position will lag behind the MEC representation, due to synaptic transmission delays (Fig. 4d). If verified, this feature of the neural population dynamics may help identify where different types of sensory inputs are integrated into the joint neural representation of position in the hippocampus and the MEC. If sensory inputs that convey spatial information project mainly to hippocampus, we expect a reversal of the time lag under conditions in which such sensory inputs are highly informative.

#### Coupling of entorhinal modules

To test whether a coupling mechanism coordinates the states of grid cell modules, we propose to simultaneously decode activities of multiple modules under conditions in which sensory inputs are absent or poor, and observe whether drifts in these modules are coordinated. A hippocampus-independent mechanism for coordinating the activity of grid cells modules, which relies on synaptic connectivity within the MEC was recently proposed (Mosheiff and Burak, 2019). Therefore, it will be of interest to further test whether the coordination requires place cell inputs to the MEC, by repeating the experiment while inactivating inputs from the hippocampus. Under these conditions, cells within a single grid cell module maintain their phase relationships (Almog et al., 2019), but it is unknown whether drifts in different modules remain coordinated. Another possible way to approach this question is to examine the coordination of grid cell modules immediately after exposure to a completely novel environment, since it is unlikely that reciprocal connectivity that could support this coordination is established immediately.

#### Variability of individual grid cell firing rates across firing fields

Two experiments can directly test the hypothesis that inputs from place cells, independent of sensory inputs, are responsible for a significant fraction of the variability observed in the firing rates of individual grid cells across different firing fields. First, if grid cell modules remain coordinated under hippocampal inactivation, it may be possible to decode position from activity in the MEC in this condition, and test whether firing rates become less variable across firing fields. Second, the independence of variability on sensory inputs could be tested by monitoring activity in the MEC under conditions in which sensory cues are poor or absent.

#### Artificial remapping

We predict that artificial remapping (Kanter et al., 2017) arises from expression of population activity patterns that mix contributions normally associated with distinct environments. To test this prediction, we propose to raise animals while exposing them to a limited number of environments. After measuring the response patterns of place cells in each of these environments, we propose to apply depolarization of the MEC in one environment, and look for overlap between place cell activity patterns, and the activity patterns that were previously observed in the other environments (with appropriate controls such as those obtained by randomly permuting cells, or by exposing the animal to new environments after the depolarization experiments). Overlap with one of the experimentally characterized spatial maps is not certain, but this is a plausible outcome of our theory, since we observe that the overlap is highly significant with many spatial maps. If successful, such experiments may provide direct evidence for a key theoretical prediction on the dynamics of auto-associative networks: the emergence of mixed states (Amit et al., 1985; Brunel, 2003).

## ACKNOWLEDGEMENTS

This research was supported by the Israel Science Foundation grant No. 1745/18, in part by the Israel Science Foundation grant No. 1978/13, and by a grant from the GIF, the German-Israeli Foundation for Scientific Research and Development. We acknowledge support from the Gatsby Charitable Foundation.

## Methods

### Place cell network

The place network consists of *N* = 4800 neurons, which represent positions in *L* environments. All environments are one-dimensional, spanning a length of 192cm with periodic boundary conditions. A distinct spatial map is generated for each environment, by choosing a random permutation that assigns all place cells to a set of preferred firing location that uniformly tile this environment.

The synaptic connectivity between place cells is expressed a sum over contributions from all spatial maps:

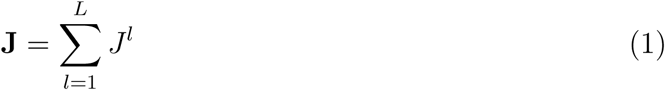

where 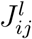 depends on the distance between the preferred firing locations of cells *i* and *j* in environment *l*, as follows

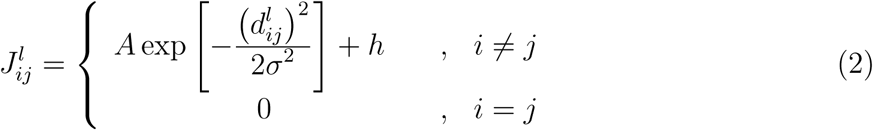

The first term is an excitatory contribution to the synaptic connectivity that decays with the distance 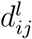, with a Gaussian profile. The second term is a uniform inhibitory contribution. The parameters *A* > 0, *σ*^2^, and *h* < 0 are listed in Table 1. Note that the connectivity matrices corresponding to any two maps *l* and *k* are related to each other by a random permutation:

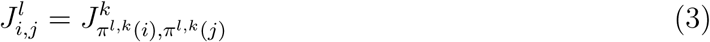

where *π*^*l,k*^ denotes the random permutation from map *k* to map *l*.

**Table 1:**

| Parameter | Value | Units |
| --- | --- | --- |
| $L$ | 6* | none |
| $A$ | $8.31 \cdot 10^{-2}$ | Hz |
| $\sigma$ | 4.8 | cm |
| $h$ | $-2.6 \cdot 10^{-2}$ | Hz |
| $B$ | $3 \cdot 10^{-3}$ | Hz |
| $\rho$ | $\frac{5}{12} \cdot 2\pi$ | radian |
| $k$ | $-2.8 \cdot 10^{-3}$ | Hz |
| $\Delta\theta$ | $\frac{5}{16} \cdot 2\pi$ | radian |
| $\alpha$ | $1.03 \cdot 10^{-2}$ | Hz |
| $\beta$ | $-6\frac{2}{3} \cdot 10^{-4}$ | Hz |
| $\tau$ | $15 \cdot 10^{-3}$ | s |
| $\Delta t$ | $2 \cdot 10^{-4}$ | s |
| $\gamma_g$ | 4 | none |
| $I_{\text{pc},0}$ | -10 | Hz |
| $\gamma_p$ | 50 | none |
| $I_{\text{gc},0}^\mu$ | $(-5, -5, -5)$ | Hz |
| $\epsilon^\mu$ | $(1.7, 1.9, 2.3)$ | $\frac{1}{\text{cm}}$ |
| $I_{\text{per}}^{\text{Depo}}$ | 500 | Hz |
| $I_{\text{per}}^{\text{Hyper}}$ | -100 | Hz |

\*In all Figures we simulated the same network with $L=6$ , except for Figure 6b and Supplementary Figure 5 where $L$ was also set to 1 and 2.

### Grid cell network

Each grid cell module is modeled as a double ring attractor [Xie et al., 2002] with *n* = 960 neurons. Within each module, neurons are assigned angles *θ*_*i*_, uniformly distributed in [0, 2*π*) (*i* = 1 … *n*). The synaptic weight between neurons *i* and *j* is given by

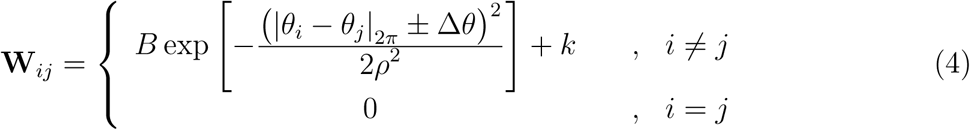

where |*θ*_*i*_ − *θ*_*j*_|_2π_ ≡ min (|*θ*_*i*_ − *θ*_*j*_|, 2π − |*θ*_*i*_ − *θ*_*j*_| and the small phase shift +Δ*θ* (−Δ*θ*) applies to the neurons with even (odd) index. Thus, evenly indexed and oddly indexed neurons comprise two sub-populations that drive motion of the activity to the right and to the left, respectively, to implement idiothetic path integration (see also Eq. 9 below). The parameters *B* > 0,*ρ*^2^, *k* < 0, and Δ*θ* are listed in Table 1. The ring attractor possesses a continuous manifold of steady states, structured as activity bumps that can be localized anywhere in the range [0, 2*π*). We refer to the the center of the activity bump as the phase of the population activity pattern. Grid cells from distinct modules are not directly connected to each other.

In our implementation of the model, we include three grid cell modules with grid spacings *λ*^*µ*^ = 38.4, 48, and 64cm for *µ* = 1, 2, and 3 respectively. Within each environment, positions are mapped to attractor states by tiling the possible phases [0, 2*π*) of the population activity periodically along the extent of the environment. For simplicity, we chose grid spacings that correspond precisely to 5, 4, and 3 cycles along the 192 cm environment. To account for grid cell realignment during global remapping [Fyhn et al., 2007], a uniform phase shift Δ^*l,µ*^ is applied to the tiling of grid phases to positions. This phase shift is randomly chosen from the range [0, 2*π*), independently for each environment *l* and module *µ*.

### Bi-directional synaptic connectivity between grid cells and place cells

The mutual connection between each place cell and grid cell depends linearly on the overlap between their tuning curves, summed over all environments. We thus define a connectivity matrix **M**^*µ*^ between all place cells and all grid cells from module *µ* as a sum over contributions *M* ^*l,µ*^ from all environments:

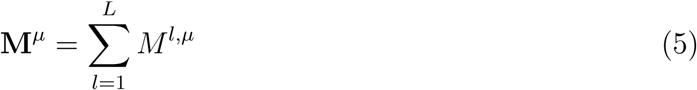

To determine *M* ^*l,µ*^ we first define a correlation matrix

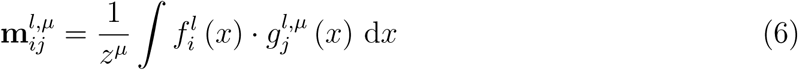

where 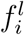 is the receptive field of place cell *i* and 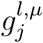is the receptive field of grid cell *j* measured from uncoupled networks at environment *l*. The normalization factor *z*^*µ*^ is chosen such that **m**^*l,µ*^ ∈ [0, 1] ⩝*µ*.

Note that the correlation matrices corresponding to any two maps *l* and *k* are related to each other by a permutation on the place cell indices, and a cyclic shift on the grid cell indices. Finally,

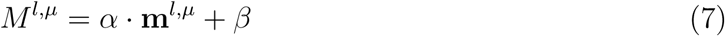

where the parameters *α* > 0 and *β* < 0 are identical for all modules and maps (Table 1).

### Dynamics

The dynamics of neural activity are described by a standard rate model. The synaptic activation *S*_*i*_ of place cell *i* evolves in time according to the following equation:

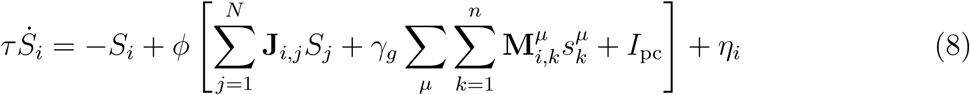

and the synaptic activation 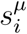 of grid cell *i* from module *µ* evolve as follows:

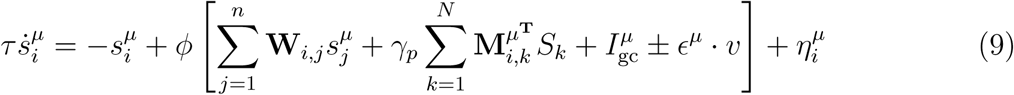

In both equations, *τ* is the synaptic time constant (taken for simplicity to be identical for all synapses). The coupling parameters *γ*_*g*_ and *γ*_*p*_ determine the strength of synaptic connections form grid cells to place cells and from place cells to grid cells, respectively. The velocity signal is denoted by *v*. The signs (+) and (-) are used for the even and odd grid cell populations, respectively. Thus, neurons with even index, whose outgoing synaptic weights are biased to the right (see Eq. 4) are preferentially activated during motion to the right, whereas neurons with odd index, whose synaptic weights are biased to the left are preferentially activated during motion to the left. The coefficients *ϵ*^*µ*^ determine the weighting of the velocity signal per module. These parameters are chosen in order to obtain correct grid spacings during idiothetic path integration. The external currents *I*_pc_ in the place cell population and 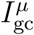 in the grid cell population are constant in time and identical in all cells from a given sub-population. These currents include two terms: one term is the baseline current, required to drive activity when a single spatial map is embedded in the connectivity. The second term compensates for interference coming from additional maps, and is proportional to *L* − 1 (see Supplementary Eqs. 36 and 37).

The transfer function *ϕ* determines the firing rate (in Hz) of place cells (*R*_*i*_) and grid cells 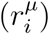 as a function of their total synaptic inputs. To resemble realistic neuronal F-I curves, it is chosen to be sub-linear:

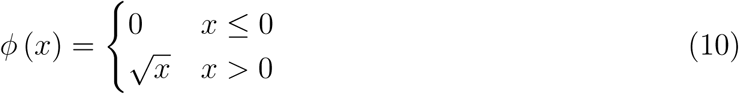

The noise terms *η*_*i*_ in Eq. 8 and 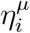 in Eq. 9 are included only in Figure 5. These terms have zero mean, and to mimic Poisson noise we assume that they are independent for different neurons and obey

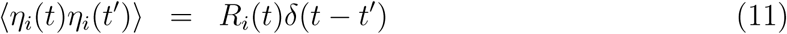

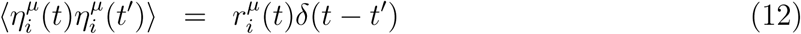

We implemented the dynamics using the Euler-method for numeric integration, with a time interval Δ*t* (Table 1).

Note that the parameter *α* in Eq. 7 is included for convenience but is redundant since it can be absorbed in the definitions of *β, γ*_*g*_, and *γ*_*p*_. We note also that due to the coupling parameters *γ*_*g*_ and *γ*_*p*_ the synaptic connectivity is not necessarily symmetric. However it is possible to recast the equations in a form that involves symmetric connectivity, by rescaling the synaptic variables *S*_*i*_ and 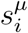 This observation is important since it implies that the dynamics always settle on stationary steady states and do not exhibit limit cycles [Cohen and Grossberg, 1983].

### Bump score and location analysis

Without loss of generality, whenever we place the system in a bump state, we do so in the map labeled as map 1.

#### Place cell network

To identify whether the place cell network expresses a bump state, and to identify its location *x* and associated spatial map *l*, we define a correlation coefficient *q*^*l*^(*x*) that quantifies the overlap between the hippocampal population activity pattern and the activity pattern corresponding to position *x* in spatial map *l*:

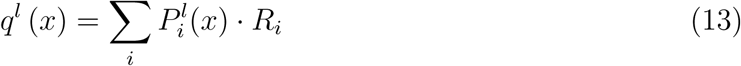

where *R*_*i*_ (already defined above) is the firing rate of place cell *i*, and 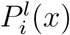, is the firing rate of neuron *i* in an idealized bump state localized at position *x* in map *l*. The idealized bump state is obtained from the activity of a network in which a single map (map *l*) is embedded in the neural connectivity, and there is no quenched noise.

Next, we define a bump score for each spatial map, defined as the maximum of the correlation coefficient *q*^*l*^(*x*) over all positions *x* in spatial map *l*:

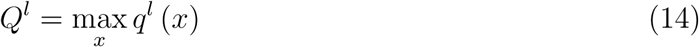

Finally, the map with the highest *Q*^*l*^ value is considered as the winning map, and the location *x* that generated that value is considered as the location of the place cell bump within that map.

#### Grid cell network

To identify the location corresponding to activity in each grid cell module, we use a similar procedure, in which we calculate the overlap between the population activity and an idealized activity pattern of grid cells, evaluated across all positions in the winning map. Note that due the periodicity of grid cell responses, multiple position produce the same correlation coefficient. When measuring the distance between the grid cell activity bump and the place activity bump or an initial (for example in Fig 3 c,g, and k), we chose from these periodically spaced positions the one closest to the location of the place cell bump.

### Grid cells hyperpolarization and depolarization implementation

To mimic to effects of transgenic DREADDs (Designer Receptors Exclusively Activated by Designer) in MEC layer2 cells as performed experimentally^49^ we added a constant current *I*_per_ to the total synaptic input driving grid cell activity. Eq. 9 is thus replaced by:

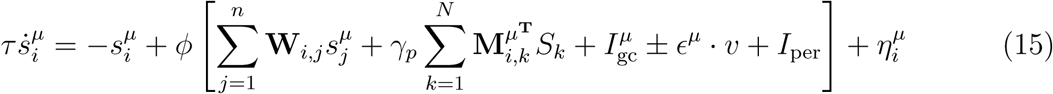

A current *I*_per_ > 0 was used for depolarization, and a current *I*_per_ < 0 was used for hyperpolarization (see Table 1).

#### Analysis of persistent states

In Figure 7, the system is placed at bump states corresponding to 500 uniformly distributed spatial locations in map 1, and its state is analyzed after a 2s delay period following the onset of grid cell depolarization.

## Supplementary Figures

**Supplementary Fig. 1:**
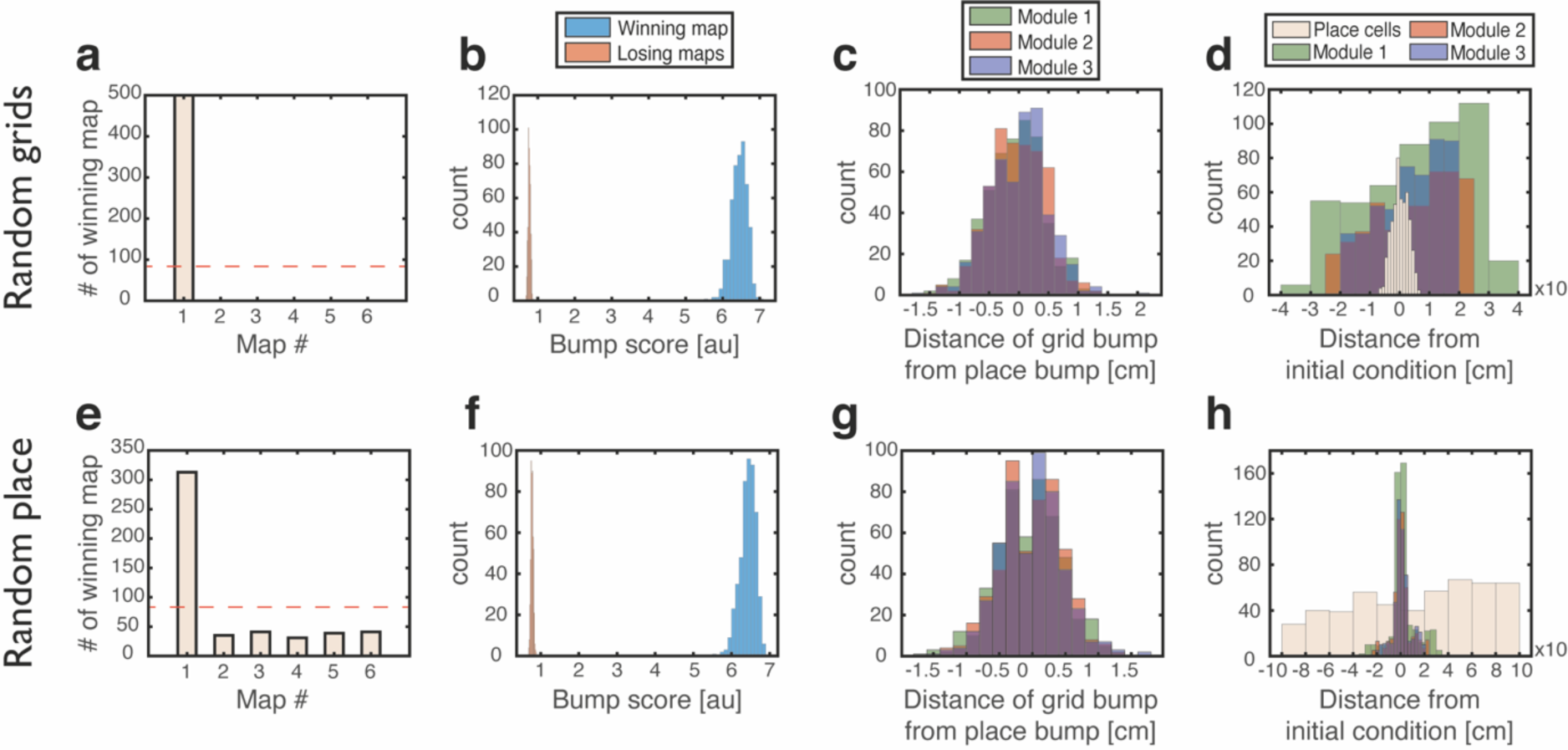
Analysis of persistent states following initial conditions of type (4). **a-d**, Similar to Figure 3a-d but for ‘Grid cells - random, Place cells - bump’ initial conditions: grid cells were initially set to have random rates, while place cells were set to encode a specific location. **e-h**, Similar to Figure 3a-d but for ‘Place cell - random, Grid cells - bump’: grid cells were set to encode a specific location, and place cells were assigned with random rates.

**Supplementary Fig. 2:**
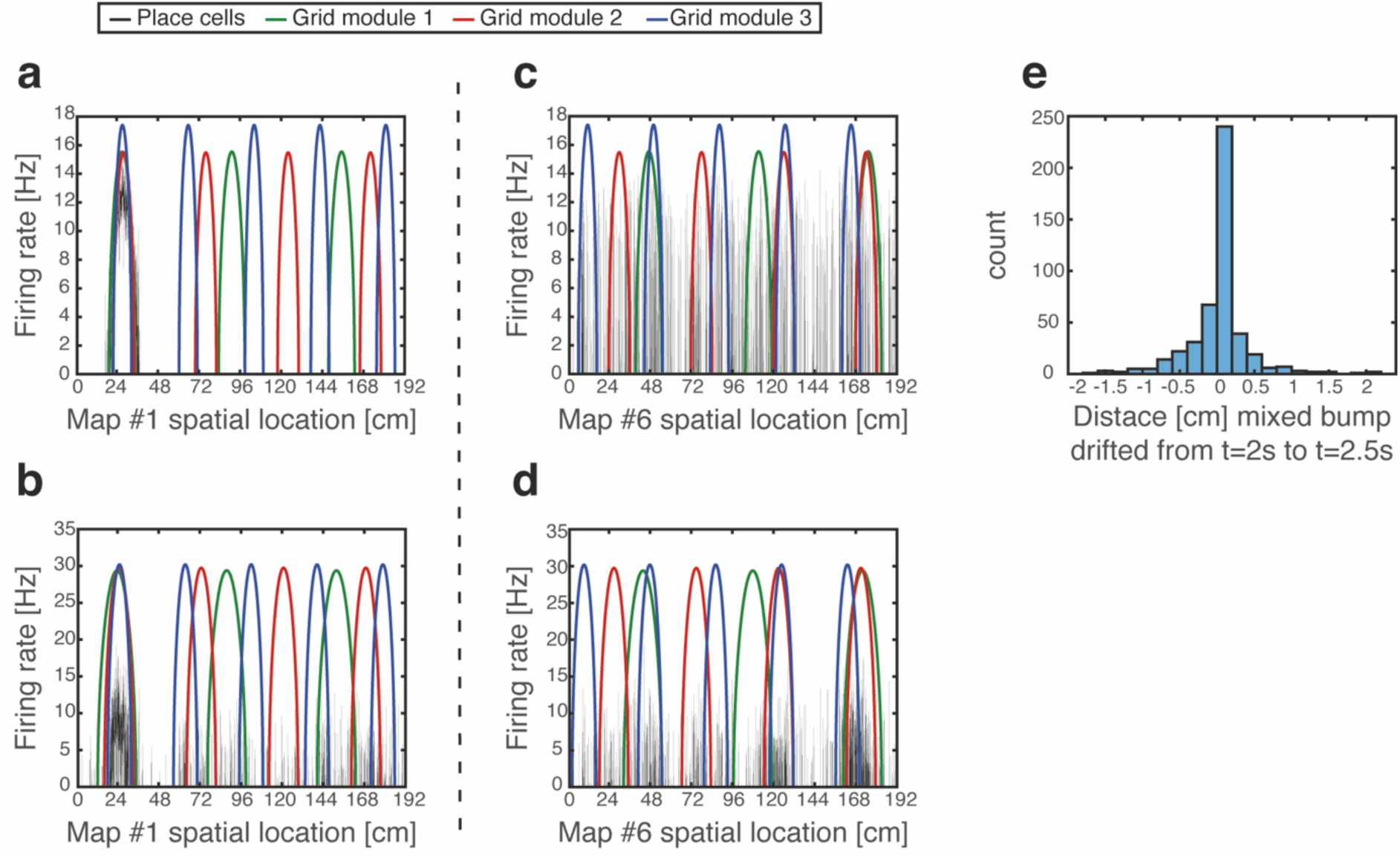
Persistent mixed state examples, and maintenance of grid cell firing location but not firing rate under grid cell depolarization. **a**, Firing rates of place cells (black) and grid cells (different modules shown in green, red and blue), shown 2.25s after starting from a ‘Consistent’ initial condition at a specific location (24cm in map #1). **b**, Same as (a) but under grid cell depolarization. Place cells exhibit scattered activity yet they preserve dense activity around their original location from (a). Note that the grid cell activity patterns continue to match the original location in map #1, but grid cells increased their firing rate. **c-d**, Same as (a-b) but observed in coordinates of map #6. Dense place cell activity can be seen around three locations (∼48cm, ∼120cm and ∼168cm) under grid cell depolarization (d). Note that these location overlap with locations in which multiple grid cell activity bumps overlap under the coordinates of map #6. **e**, Analysis of persistence in the mixed state. Location of activity place cell bumps of winning maps was measured using the bump score as in Figure 3 (Methods), 2s and 2.5s after onset of depolarization. A histogram of the changes in the measured location within this period is shown, demonstrating that drifts in persistent mixed states are small. Out of 500 simulations, 29 simulations in which the identity of the winning map changed within this period are excluded from the histogram.

**Supplementary Fig. 3:**
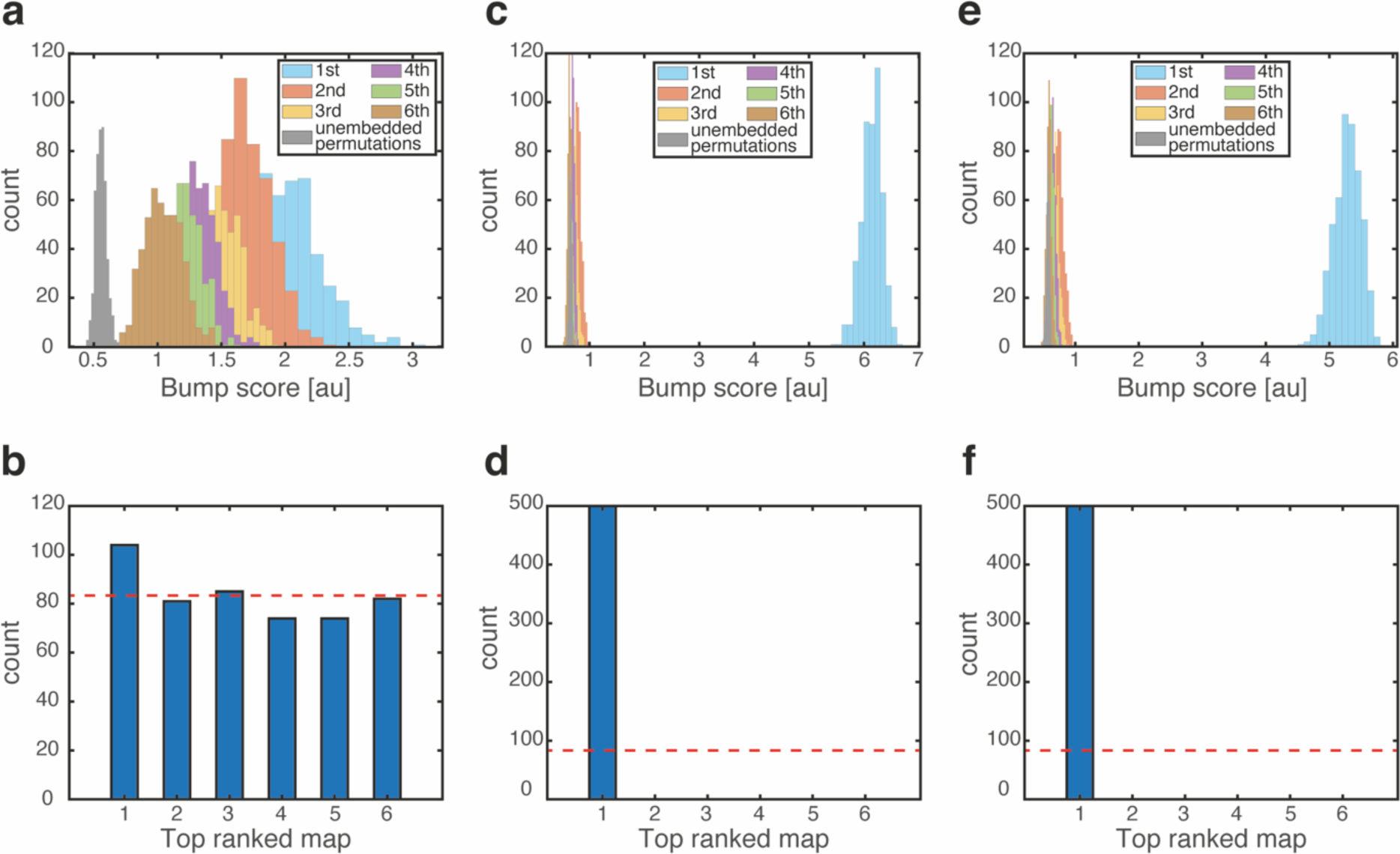
Mixed states can be induced or suppressed by changing the connectivity between grid cells and place cells. **a-b**, Same analysis as Figure 7a-b but while doubling the bi-directional synaptic strengths between grid cells and place cells. Mixed states emerge under this manipulation, without grid cell depolarization. **c-d**, Same as (a-b) but while selectively enhancing only the contribution to the connectivity associated with map #1. Mixed states do not emerge in this case. **e-f**, Same as (c-d) but under grid cell depolarization. Although winning map bump score (cyan) is smaller than in Figure 3b mixed states still do not emerge in this case.

**Supplementary Fig. 4:**
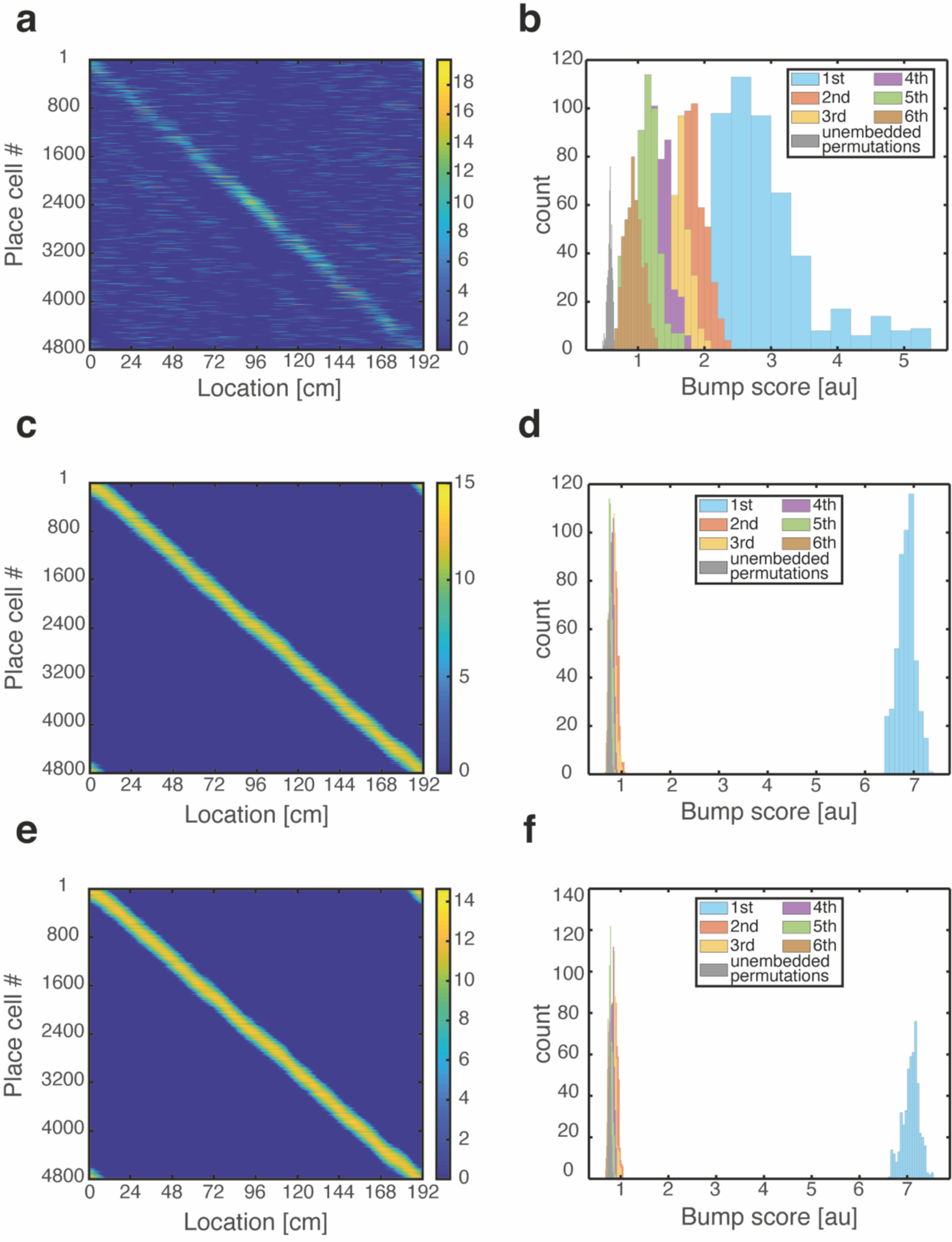
Place cell population activity patterns observed while traversing the environment using path integration. **a**, Place cell firing rates vs. spatial location under grid cell depolarization and grid cell integration of velocity input that produces one complete cycle through the environment. Color bar: place cell firing rates [Hz]. Place cells exhibit unimodal and continuous firing fields. Same result is obtained while traversing the environment with different velocities, direction, or number of cycles through the environment (not shown). **b**, Similar analysis as in Figure 7b, showing the existence of mixed states throughout the process. Activity patterns were analyzed at 500 time points uniformly distributed across the cycle. **c-d**, Same as (a-b) but without grid cell depolarization (control). **e-f**, Same as (a-b) during grid cell hyperpolarization.

**Supplementary Fig. 5:**
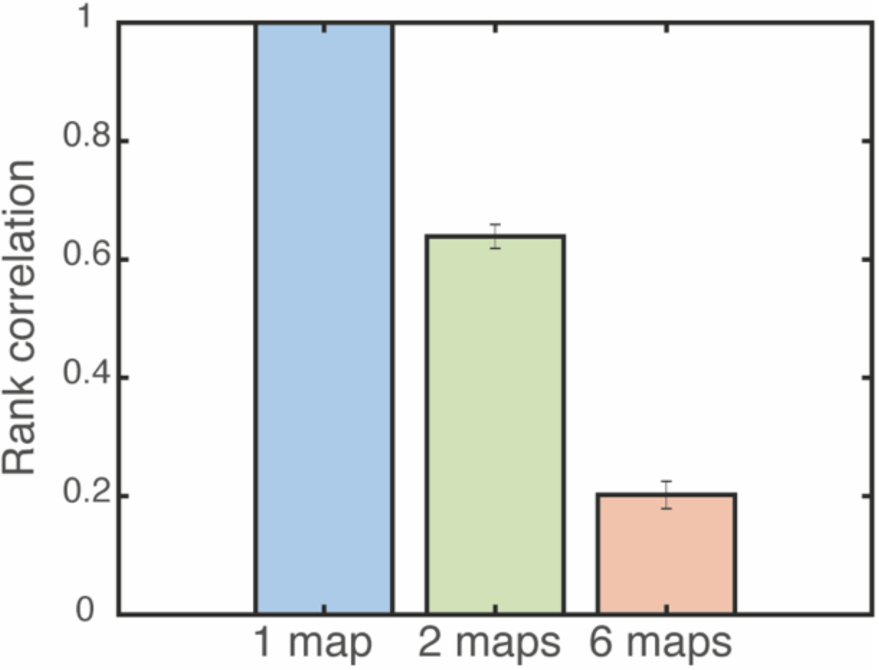
Altered firing rate relationship between individual grid fields is observed during grid cell depolarization. Spearman rank correlation before and after grid cell depolarization is shown for a varied number of embedded maps. Error bars are 1.96 times the standard deviations obtained from each simulation, divided by the square root of the number of realizations (corresponding to a confidence interval of 95%).

## Supplementary information

In this supplementary text we analyze mathematically how the embedding of multiple spatial maps in the synaptic connectivity affects the existence of bump states in the system, using a mean-field analysis. Similar questions have been extensively studied for associative networks with discrete memories, and for models of multiple spatial map representations in the hippocampus. Here we consider properties of the coupled entorhinal-hippocampal network.

As the number of spatial maps *L* increases, the shape of the bump is increasingly distorted until at a certain critical *L*_*c*_ the representation of the bump will totally collapse and no localized bump will be observed in any of the embedded maps. This happens since with every map added, there is addition of quenched noise into the place cell connections, arising from the random permutations and thus, there is accumulating interference between the distinct map representations. To characterize the influence of the quenched noise, we compute here the mean and variance of quenched noisy inputs to place cells and grid cells.

The calculation of the mean input is used to adjust the feed-forward inhibition in the network (which is uniform in each grid cell module and within the place cell network), such that the mean input to all cells remains fixed with addition of new maps (Eqs. 36 and 37). The calculation of the variance allows us to conclude that *L*_*c*_ is proportinal to the number of neurons in the network, when the fraction of active cells and their typical firing rates are kept fixed with addition of new spatial maps.

### Quenched noise conveyed to place cells

Without loss of generality, we assume a steady bump state in map 1. Since the synaptic activities are steady, we can replace them in all expressions with the corresponding firing rates: *S*_*i*_ = *R*_*i*_ and *s*_*i*_ = *r*_*i*_. The firing rate profile of place cells can be thought as the sum of two contributions: a smooth bump of synaptic activation associated with map 1, and noisy deviations originating from the synaptic weights associated with the rest of the maps. When *L* ≥ *L*_*c*_ these deviations are too large and the representation collapses.

Since we assume that the deviations are sufficiently small, we decompose terms in the steady state equation to inputs from map 1 and to inputs from all of the other interfering maps. We obtain terms describing (1) inputs coming from place cells through the synaptic connectivity that corresponds to map 1, (2) noisy inputs coming from place cells arising from all the other interfering maps, (3) inputs coming from grid cells, associated with map 1, and (4) noisy inputs coming from grid cells that arise from all the other interfering maps. Rewriting the steady state equation of a place cell accordingly yields

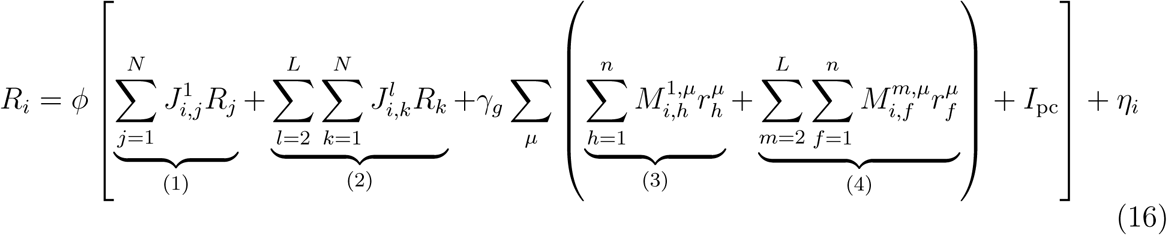

We are interested in the means and variances of terms (2) and (4), compared to terms (1) and (3).

### Quenched noise conveyed to place cells by place cells (term 2)

We note that the sum of each row and the sum of each column of *J* ^*l*^ are all identical, and denote

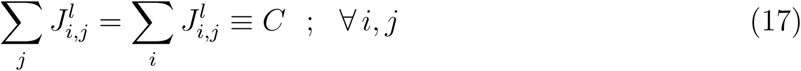

We note also that ⟨*J*^*l*^*J*^*m*^⟩ = ⟨*J*^*l*^⟩ ⟨*J*^*m*^⟩ for *l* ≠ where the pointy brackets denote an average over the random permutation *π*^*l*^ that relates the matrix *J* ^*l*^ to *J* ^1^:

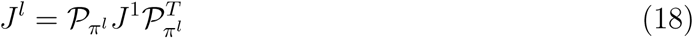

and 𝒫_*π*_ is the permutation matrix corresponding to the permuation *π*.

For convenience, we denote term (2) by *P* :

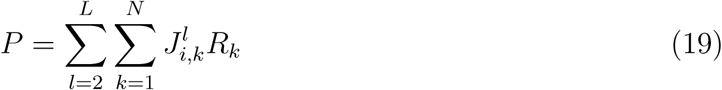

The mean of *P* is

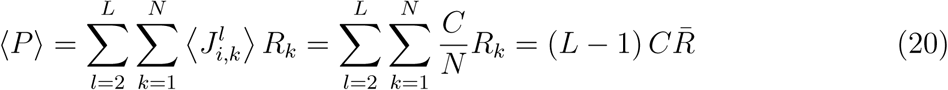

where 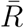 is the average firing rate of place cells in the bump steady state when *L* = 1. Since the permutations in different maps are drawn independently, the variance of *P* is

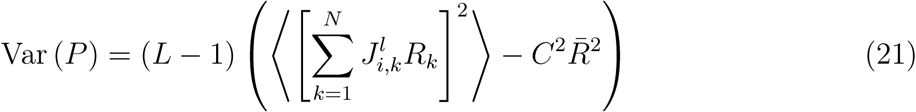

Thus, both the mean and the variance grow linearly with *L*, and vanish when *L* = 1.

### Quenched noise conveyed to place cells by grid cells (term 4)

Similarly, the sums of all rows of **m**^*l,µ*^ are equal to each other, and we denote

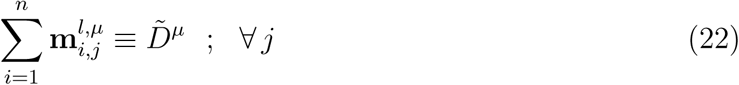

Thus,

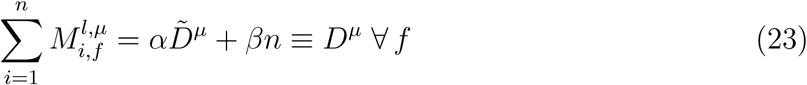

Note also that ⟨*M*^*l,μ*^*M*^*m,μ*^⟩ = ⟨*M*^*l,μ*^⟩ ⟨*M*^*m,μ*^⟩ for all *l* ≠ *m*.

For convenience, we denote the contribution to term 4 from module *µ* by *G*^*µ*^:

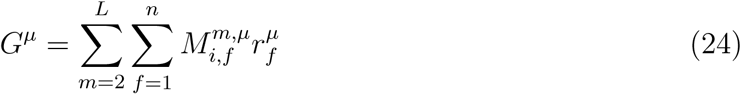

The mean contribution of module *µ* to the quenched noise in the input to place cells is

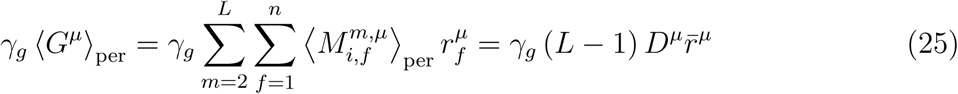

and

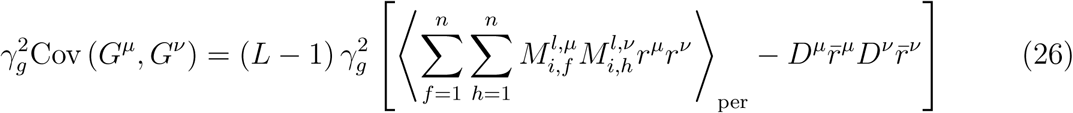

where 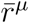 is the average firing rate of grid cells from module *µ* in the bump steady state when *L* = 1, and using the independence of permutations in different maps. Again, we see the same linear dependence of the mean and variance with *L* as seen by the inputs conveyed by place cells.

### Covariance of place cell and grid cell inputs

Similarly,

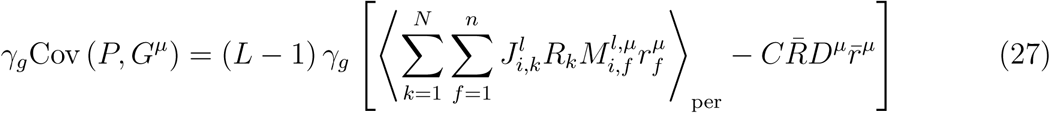

### Overall mean and variance

The overall mean of the quenched noise contribution to the place cell inputs is:

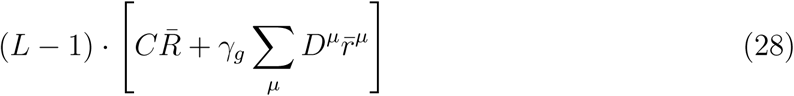

and the variance is:

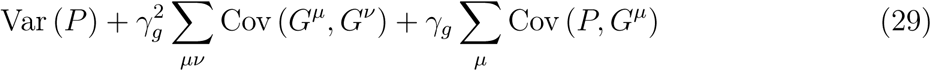

Note that all terms in this expressions are proportional to (L − 1).

### Quenched noise conveyed to grid cells

Similarly to our derivation for place cells, we decompose the input to grid cells into a contribution from map 1 (in which we assume the bump state), and to inputs from all other interfering maps:

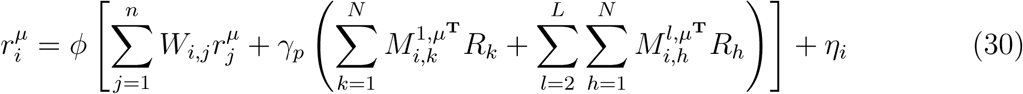

The quenched noise inputs to grid cells originate solely from place cells. Thus, we are interested in the mean and variance of the third term, compared to the sum of the first two terms in the argument of *ϕ*. We denote

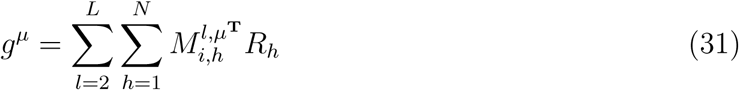

We note that

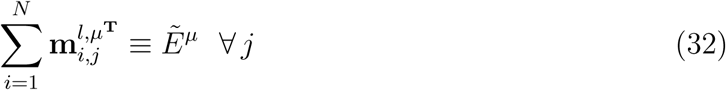

and thus

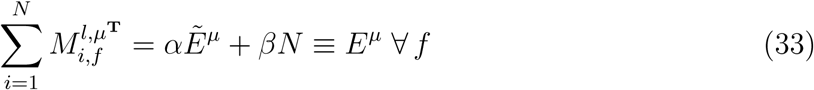

From here on the derivation is similar to the previous calculation and it is straightforward to see that the mean of the quenched noise input to grid cell is equal to

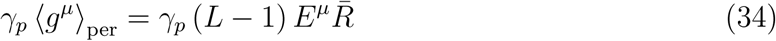

and the variance is

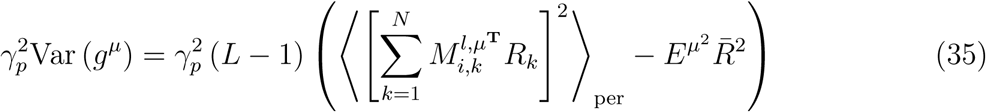

### Compensating for the mean of the quenched noise

In order to sustain a constant mean input to place cells and grid cells as *L* is varied, we adjust the external current to compensate for the mean of the quenched noise, Eqs. 28 and 34. Thus, the external input to place cells is

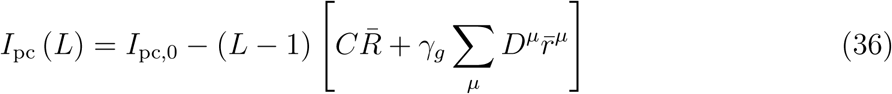

and the external input to grid cells is

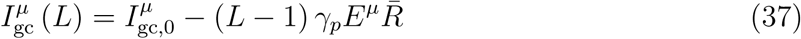

where *I*_pc,0_ and 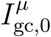 are independent of *L*.

### Scaling of *L*_*c*_ with *N*

We next consider how the variance of inputs scales with the number of neurons in the system. for simplicity, we consider scaling of all neuron population sizes by a common pre-factor *a*:

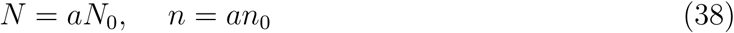

and

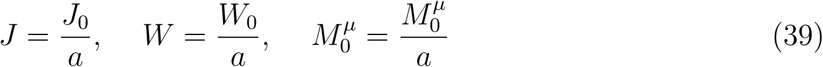

Under the assumption that *N* >> 1 place cells cover uniformly and densely the environment, the distribution of place cell firing rates at steady state remains fixed under re-scaling.

The signal conveyed to place cells in Eq.(16), denoted by *S*_0_, is composed of the contribution of terms 1 and 3. The noise conveyed to place cells, denoted by 𝒩_0_, is composed of the contribution of terms 2 and 4. It is straightforward to show that after re-scaling the system

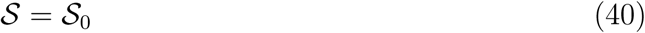

The mean of the noise is canceled by the adjustment of the external current (Eq.36),

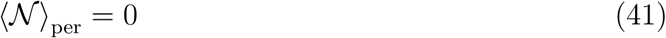

However,

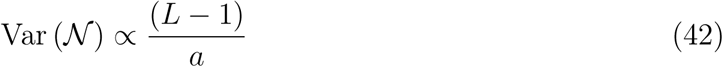

Therefore, as the number of place cells *N* increases, the number of possible embedded maps *L*_*c*_ also increases.

## REFERENCES

de Almeida, L., Idiart, M., and Lisman, J.E. (2009). The input–output transformation of the hippocampal granule cells: from grid cells to place fields. J. Neurosci. 29, 7504–7512.

Almog, N., Tocker, G., Bonnevie, T., Moser, E.I., Moser, M.B., and Derdikman, D. (2019). During hippocampal inactivation, grid cells maintain synchrony, even when the grid pattern is lost. Elife 8.

Amit, D.J., Gutfreund, H., and Sompolinsky, H. (1985). Spin-glass models of neural networks. Phys. Rev. A 32, 1007.

Battaglia, F.P., and Treves, A. (1998). Attractor neural networks storing multiple space representations: A model for hippocampal place fields. Phys. Rev. E 58, 7738–7753.

Ben-Yishai, R., Bar-Or, R.L., and Sompolinsky, H. (1995). Theory of orientation tuning in visual cortex. Proc. Natl. Acad. Sci. 92, 3844–3848.

Brandon, M.P., Koenig, J., Leutgeb, J.K., and Leutgeb, S. (2014). New and Distinct Hippocampal Place Codes Are Generated in a New Environment during Septal Inactivation. Neuron 82, 789–796.

Brunel, N. (2003). Course 10 Network models of memory. In Les Houches Summer School Proceedings, p. 407.

Burak, Y. (2014). Spatial coding and attractor dynamics of grid cells in the entorhinal cortex. Curr. Opin. Neurobiol. 25, 169–175.

Burak, Y., and Fiete, I.R. (2009). Accurate Path Integration in Continuous Attractor Network Models of Grid Cells. PLoS Comput. Biol. 5, e1000291.

Cohen, M.A., and Grossberg, S. (1983). Absolute Stability of Global Pattern Formation and Parallel Memory Storage by Competitive Neural Networks. IEEE Trans. Syst. Man Cybern. 13, 815–826.

D’Albis, T., and Kempter, R. (2017). A single-cell spiking model for the origin of grid-cell patterns. PLoS Comput. Biol. 13, e1005782.

Deadwyler, S.A., West, J.R., Cotman, C.W., and Lynch, G. (1975). Physiological studies of the reciprocal connections between the hippocampus and entorhinal cortex. Exp. Neurol. 49, 35–57.

Dordek, Y., Soudry, D., Meir, R., and Derdikman, D. (2016). Extracting grid cell characteristics from place cell inputs using non-negative principal component analysis. Elife 5, e10094.

Dunn, B., Wennberg, D., Huang, Z., and Roudi, Y. (2017). Grid cells show field-to-field variability and this explains the aperiodic response of inhibitory interneurons. ArXiv Prepr. ArXiv 1701, 04893.

Fiete, I.R., Burak, Y., and Brookings, T. (2008). What grid cells convey about rat location. J. Neurosci. 28, 6858–6871.

Fuhs, M.C., and Touretzky, D.S. (2006). A spin glass model of path integration in rat medial entorhinal cortex. J. Neurosci. 26, 4266–4276.

Fyhn, M., Hafting, T., Treves, A., Moser, M.B., and Moser, E.I. (2007). Hippocampal remapping and grid realignment in entorhinal cortex. Nature 446, 190–194.

Gardner, R.J., Lu, L., Wernle, T., Moser, M.B., and Moser, E.I. (2019). Correlation structure of grid cells is preserved during sleep. Nat. Neurosci. 22, 598–608.

Gil, M., Ancau, M., Schlesiger, M.I., Neitz, A., Allen, K., De Marco, R.J., and Monyer, H. (2018). Impaired path integration in mice with disrupted grid cell firing. Nat. Neurosci. 21, 81–91.

Guanella, A., Kiper, D., and Verschure, P. (2007). A model of grid cells based on a twisted torus topology. Nternational J. Neural Syst. 17, 231–240.

Hafting, T., Fyhn, M., Molden, S., Moser, M.B., and Moser, E.I. (2005). Microstructure of a spatial map in the entorhinal cortex. Nature 436, 801–806.

Hales, J.B., Schlesiger, M.I., Leutgeb, J.K., Squire, L.R., Leutgeb, S., and Clark, R.E. (2014). Medial entorhinal cortex lesions only partially disrupt hippocampal place cells and hippocampus-dependent place memory. Cell Rep. 9, 893–901.

Hopfield, J.J. (1982). Neural networks and physical systems with emergent collective computational abilities. Proc. Natl. Acad. Sci. 79, 2554–2558.

Ismakov, R., Barak, O., Jeffery, K., and Derdikman, D. (2017). Grid cells encode local positional information. Curr. Biol. 27, 2337–2343.

Itskov, V., Hansel, D., and Tsodyks, M. (2011). Short-term facilitation may stabilize parametric working memory trace. Front. Comput. Neurosci. 5, 40.

Jeffery, K.J. (2011). Place cells, grid cells, attractors, and remapping. Neural Plast.

Jun, J.J., Steinmetz, N.A., Siegle, J.H., Denman, D.J., Bauza, M., Barbarits, B., Lee, A.K., Anastassiou, C.A., Andrei, A., Aydin, Ç., et al. (2017). Fully integrated silicon probes for high-density recording of neural activity. Nature 551, 232–236.

Kanter, B.R., Lykken, C.M., Avesar, D., Weible, A., Dickinson, J., Dunn, B., Borgesius, N.Z., Roudi, Y., and Kentros, C.G. (2017). A Novel Mechanism for the Grid-to-Place Cell Transformation Revealed by Transgenic Depolarization of Medial Entorhinal Cortex Layer II. Neuron 93, 1480-1492.e6.

Koenig, J., Linder, A.N., Leutgeb, J.K., and Leutgeb, S. (2011). The spatial periodicity of grid cells is not sustained during reduced theta oscillations. Science (80-.). 332, 592–595.

Kropff, E., and Treves, A. (2008). The emergence of grid cells: Intelligent design or just adaptation? Hippocampus 18, 1256–1269.

Kropff, E., Carmichael, J.E., Moser, M.B., and Moser, E.I. (2015). Speed cells in the medial entorhinal cortex. Nature 523, 419–424.

Langston, R.F., Ainge, J.A., Couey, J.J., Canto, C.B., Bjerknes, T.L., Witter, M.P., Moser, E.I., and Moser, M.B. (2010). Development of the spatial representation system in the rat. Science (80-.). 328, 1576–1580.

Laptev, D., and Burgess, N. (2019). Neural Dynamics Indicate Parallel Integration of Environmental and Self-Motion Information by Place and Grid Cells. Front. Neural Circuits 13, 59.

Mathis, A., Herz, A. V., and Stemmler, M.B. (2012). Resolution of nested neuronal representations can be exponential in the number of neurons. Phys. Rev. Lett. 109, 018103.

McNaughton, B.L., Barnes, C.A., Gerrard, J.L., Gothard, K., Jung, M.W., Knierim, J.J., Kudrimoti, H., and Qin, Y. (1996). Deciphering the hippocampal polyglot: the hippocampus as a path integration system. J. Exp. Biol. 199, 173–185.

McNaughton, B.L., Battaglia, F. P. J., Ensen, O., Moser, E.I., and Moser, M.B. (2006). Path integration and the neural basis of the “cognitive map.” Nat. Rev. Neurosci. 7, 663–678.

Miao, C., Cao, Q., Ito, H.T., Yamahachi, H., Witter, M.P., Moser, M.B., and Moser, E.I. (2015). Hippocampal Remapping after Partial Inactivation of the Medial Entorhinal Cortex. Neuron 88, 590–603.

Monaco, J.D., and Abbott, L.F. (2011). Modular realignment of entorhinal grid cell activity as a basis for hippocampal remapping. J. Neurosci. 31, 9414–9425.

Monasson, R., and Rosay, S. (2013). Crosstalk and transitions between multiple spatial maps in an attractor neural network model of the hippocampus: Phase diagram. Phys. Rev. E - Stat. Nonlinear, Soft Matter Phys. 87, 062813.

Monasson, R., and Rosay, S. (2014). Crosstalk and transitions between multiple spatial maps in an attractor neural network model of the hippocampus: Collective motion of the activity. Phys. Rev. E 89, 032803.

Monsalve-Mercado, M.M., and Leibold, C. (2017). Hippocampal Spike-Timing Correlations Lead to Hexagonal Grid Fields. Phys. Rev. Lett. 119, 038101.

Mosheiff, N., and Burak, Y. (2019). Velocity coupling of grid cell modules enables stable embedding of a low dimensional variable in a high dimensional neural attractor. Elife 8.

Muller, R.U., and Kubie, J.L. (1987). The effects of changes in the environment on the spatial firing of hippocampal complex-spike cells. J. Neurosci. 7, 1951–1968.

Neher, T., Azizi, A.H., and Cheng, S. (2017). From grid cells to place cells with realistic field sizes. PLoS One 12.

O’Keefe, J., and J., D. (1971). The hippocampus as a spatial map: preliminary evidence from unit activity in the freely-moving rat. Brain Res. 34, 171–175.

Pfeiffer, B.E., and Foster, D.J. (2015). Autoassociative dynamics in the generation of sequences of hippocampal place cells. Science (80-.). 349, 180–183.

Quirk, G.J., Muller, R.U., and Kubie, J.L. (1990). The firing of hippocampal place cells in the dark reflects the rat’s recent experience. J. Neurosci. 10, 2008–2017.

Redish, A.D., Elga, A.N., and Touretzky, D.S. (1996). A coupled attractor model of the rodent Head Direction system. Netw. Comput. Neural Syst. 7, 671–685.

Rennó-Costa, C., and Tort, A.B. (2017). Place and grid cells in a loop: implications for memory function and spatial coding. J. Neurosci. 37, 8062–8076.

Rolls, E.T., Stringer, S.M., and Elliot, T. (2006). Entorhinal cortex grid cells can map to hippocampal place cells by competitive learning. Netw. Comput. Neural Syst. 17, 447–465.

Samsonovich, A., and McNaughton, B.L. (1997). Path integration and cognitive mapping in a continuous attractor neural network model. J. Neurosci. 17, 5900–5920.

Sargolini, F., Fyhn, M., Hafting, T., McNaughton, B.L., Witter, M.P., Moser, M.B., and Moser, E.I. (2006). Conjunctive representation of position, direction, and velocity in entorhinal cortex. Science (80-.). 312, 758–762.

Schlesiger, M.I., Boublil, B.L., Hales, J.B., Leutgeb, J.K., and Leutgeb, S. (2018). Hippocampal Global Remapping Can Occur without Input from the Medial Entorhinal Cortex. Cell Rep. 22, 3152–3159.

Skaggs, W.E., Knierim, J.J., Kudrimoti, H.S., and McNaughton, B.L. (1995). A model of the neural basis of the rat’s sense of direction. Adv. Neural Inf. Process. Syst. 173–180.

Solstad, T., Moser, E.I., and Einevoll, G.T. (2006). From grid cells to place cells: A mathematical model. Hippocampus 16, 1026–1031.

Sreenivasan, S., and Fiete, I.R. (2011). Grid cells generate an analog error-correcting code for singularly precise neural computation. Nat. Neurosci. 14, 1330.

Stensola, H., Stensola, T., Solstad, T., Frøland, K., Moser, M.B., and Moser, E.I. (2012). The entorhinal grid map is discretized. Nature 492, 72–78.

Stepanyuk, A. (2015). Self-organization of grid fields under supervision of place cells in the model of neuron with associative plasticity. Biol. Inspired Cogn. Archit. 13, 48–62.

Van Strien, N.M., Cappaert, N.L.M., and Witter, M.P. (2009). The anatomy of memory: An interactive overview of the parahippocampal-hippocampal network. Nat. Rev. Neurosci. 10, 272–282.

Trettel, S.G., Trimper, J.B., Hwaun, E., Fiete, I.R., and Colgin, L.L. (2019). Grid cell co-activity patterns during sleep reflect spatial overlap of grid fields during active behaviors. Nat. Neurosci. 22, 609–617.

Weber, S.N., and Sprekeler, H. (2018). Learning place cells, grid cells and invariances with excitatory and inhibitory plasticity. Elife 7, e34560.

Wei, X.X., Prentice, J., and Balasubramanian, V. (2015). A principle of economy predicts the functional architecture of grid cells. Elife 4, e08362.

Welinder, P.E., Burak, Y., and Fiete, I.R. (2008). Grid cells: The position code, neural network models of activity, and the problem of learning. Hippocampus 18, 1283–1300.

Wills, T.J., Lever, C., Cacucci, F., Burgess, N., and O’Keefe, J. (2005). Attractor dynamics in the hippocampal representation of the local environment. Science 308, 873–876.

Wills, T.J., Cacucci, F., Burgess, N., and O’Keefe, J. (2010). Development of the hippocampal cognitive map in preweanling rats. Science (80-.). 328, 1573–1576.

Xie, X., Hahnloser, R.H.R., and Seung, H.S. (2002). Double-ring network model of the head-direction system. Phys. Rev. E - Stat. Physics, Plasmas, Fluids, Relat. Interdiscip. Top. 66, 041902.

Yoon, K., Buice, M.A., Barry, C., Hayman, R., Burgess, N., and Fiete, I.R. (2013). Specific evidence of low-dimensional continuous attractor dynamics in grid cells. Nat. Neurosci. 16, 1077.

Zhang, K. (1996). Representation of spatial orientation by the intrinsic dynamics of the head-direction cell ensemble: A theory. J. Neurosci. 16, 2112–2126.

Zhang, S.J., Ye, J., Miao, C., Tsao, A., Cerniauskas, I., Ledergerber, D., Moser, M.B., and Moser, E.I. (2013). Optogenetic dissection of entorhinal-hippocampal functional connectivity. Science (80-.). 340, 1232627.

Ziv, Y., Burns, L.D., Cocker, E.D., Hamel, E.O., Ghosh, K.K., Kitch, L.J., El Gamal, A., and Schnitzer, M.J. (2013). Long-term dynamics of CA1 hippocampal place codes. Nat. Neurosci. 16, 264.

Zong, W., Wu, R., Li, M., Hu, Y., Li, Y., Li, J., Rong, H., Wu, H., Xu, Y., Lu, Y., et al. (2017). Fast high-resolution miniature two-photon microscopy for brain imaging in freely behaving mice. Nat. Methods 14, 713–719.

